# Dynamic regulation of H2A.Zub and H3K27me3 by ambient temperature in plant cell fate determination

**DOI:** 10.1101/2024.07.18.604029

**Authors:** Kehui Zhu, Long Zhao, Fangfang Lu, Xuelei Lin, Chongsheng He, Doris Wagner, Jun Xiao

## Abstract

Crucial to plant development, ambient temperature triggers intricate mechanisms enabling adaptive responses to temperature variations. The precise coordination of chromatin modifications in shaping cell fate under diverse temperatures remains elusive. Our study, integrating comprehensive transcriptome, epigenome profiling and genetics, unveils that lower ambient temperature (16°C) restores developmental defects caused by H3K27me3 loss in PRC2 mutants by specifically depositing H2A.Zub at ectopically expressed embryonic genes, such as *ABI3* and *LEC1*. This deposition leads to re-silencing of these genes and compensates for H3K27me3 depletion. PRC1-mediated H2A.Zub and PRC2-catalyzed H3K27me3 play roles in silencing transcription of these embryonic genes for post-germination development. Low temperature decelerates H2A.Z turnover at specific loci likely by attenuating the interaction between TOE1 and H2A.Z chaperone, sustaining repression of embryonic genes and alleviating requirement for PRC2-H3K27me3 at post-germination stage. Our findings offer mechanistic insights into the cooperative epigenetic layers facilitating plants adaptation to varying environmental temperatures.

## Introduction

The dynamic development of plants is influenced by both genetic information and external environmental cues. Epigenetic regulation, including DNA methylation, histone modification, and chromosome remodeling, collectively shapes gene expression by modifying chromatin structure and gene accessibility, thereby impacting plant growth and development.^1,2^ Environmental factors like light, temperature, water, and nutrients are crucial for plant growth, especially temperature, which affects growth rate, seed germination, and flowering time.^3–5^ The intricate interplay between epigenetic regulation and environmental cues allows plants to adapt their developmental processes to various conditions.

Polycomb group (PcG) proteins are conserved and crucial epigenetic regulators involved in determining plant developmental fate,^6,7^ which comprising Polycomb repressive complex 1 (PRC1) that mediate H2Aub or H2A.Zub and PRC2 that mediate H3K27me3.^8–10^ These complexes establish inhibitory chromatin states, crucial for spatio-temporal specific gene expression regulation during various developmental transitions and cell fate determining.^6,11–13^ Thus, mutations in PRC1 or PRC2 lead to diverse developmental defects like abnormal cotyledons, root swelling, early flowering, and seed abortion.^14–17^ Notably, embryo-to-seedling transition exemplifies the cooperation of PRC1 and PRC2.^11,14^ PRC1 initiates repression of seed maturation genes and embryonic genes, such as *ABA INSENSITIVE 3* (*ABI3*)*, LEAFY COTYLEDON 1* (*LEC1*)*, LEC2, FUSCA3* (*FUS3*) and *BABY BOOM* (*BBM*), ensuring proper germination.^14,18,19^ PRC2 then maintains this inhibition through H3K27me3, promoting normal seedling development. Therefore, mutations in PRC1 core components, like double mutant of *ATRING1A* and *ATRING1B* (*Atring1a Atring1b*), show embryonic traits at cotyledons and swollen primary root post-germination,^11,14^ while mutations in PRC2 core components, like double mutant of *CURLY LEAF* (*CLF*) and *SWINGER* (*SWN*) (*clf-28 swn-7*), lead to callus-like undifferentiated cells until cotyledon fully opening.^14,17^ The current understanding of PRC1 and PRC2 functions in the post-germination stage is primarily based on phenotypic evidence from mutants and changes in H2Aub and H3K27me3 levels on a few specific genes at isolated time points. However, dynamic changes in histone modifications at the genomic level across different developmental stages after germination have not been thoroughly tracked.

Plants can perceive subtle temperature changes and adjust their development. Higher temperatures induce thermomorphogenesis, characterized by elongated hypocotyls, petioles, and leaf hyponasty,^20,21^ while lower temperatures slow growth, leading to delayed flowering.^22^ Epigenetic regulation plays a crucial role in this temperature-responsive developmental process. Currently, H2A.Z, a highly conserved histone variant that can modulate chromatin structure and gene expression,^23–25^ has been reported to sense temperature changes across eukaryotes.^26^ H2A.Z establishes low gene accessibility at +1 nucleosome and high gene accessibility at −1 nucleosome. In addition, H2A.Z’s impact on transcription is influenced by post-translational modifications, for example, acetylated H2A.Z activates target genes, while monoubiquitinated H2A.Z represses transcription.^9,27^ In *Arabidopsis*, the chromatin remodeling complexes Inositol Requiring 80 (INO80) and SWi2/snf2-Related 1 (SWR1) complex (INO80-C and SWR1-C) regulate the incorporation of H2A.Z into nucleosomes.^28,29^ INO80-C has been reported to connect with H2A.Z eviction and active expression at a subset of genes to promote thermomorphogenesis. It can directly interact with transcription factor PHYTOCHROME INTERACTING FACTOR 4 (PIF4) to target specific genes.^30^ At higher temperature, H2A.Z is removed from auxin-related genes like *INDOLE-3-ACETIC ACID INDUCIBLE 29* (*IAA29*) and *YUCCA8* (*YUC8*), facilitating hypocotyl elongation and from *SOMNUS* (*SOM*), inhibiting seed germination.^30–32^ Conversely, low temperatures increase H2A.Z occupancy at Heat Shock Transcription Factor (HSF)–Heat Shock Protein (HSP) pathway genes and at *INDEHISCENT* (*IND*) to delay fruit dehiscence in *Arabidopsis*.^26,33,34^ These studies highlight the crucial role of H2A.Z in plant response to temperature changes. However, the mechanisms underlying plant responses to long-term environmental temperature changes, and their complex effects on plant development, remain poorly understood.

The interplay of epigenetic modifications is crucial for cell fate development and environmental responsiveness by shaping gene accessibility and transcriptional regulation. For instance, PRC1 and PRC2 exhibit intricate relationship: loss of B cell-specific Moloney murine leukemia virus Integration site 1 (AtBMI1) activity in *atbmi1a^−/−^b^−/−^c^+/^* results in a general decrease of H3K27me3^35^ but in *Atbmi1a-1 Atbmi1b* intermediate mutants H3K27me3 level increases at seed maturation genes and stem cell regulators, whereas a lack of H3K27me3 leads to increased H2Aub at these genes.^14^ The detailed crosstalk between PRC1 and PRC2 remains incompletely understood. H2A.Z can co-localize with H3K4me3 or H3K27me3 to regulate gene expression.^24^ Besides, H2A.Z eviction at high temperature may be companied by H3K4me3 deposition or H3K9 deacetylation to promote thermomorphogenesis.^30–32^ This intricate interplay between environmental cues and epigenetic regulation empowers plants to adaptively adjust their developmental trajectories in diverse growing conditions.

In this study, we explore how low ambient temperature and chromatin regulation intersect to influence cell fate using *prc2* mutants, which exhibit a callus-like morphology. We find that under low temperature, these mutants show specific accumulation of H2A.Z at embryonic-related genes without H3K27me3, restoring proper cell fate. Unlike at 22°C, where H2A.Z levels decrease after cotyledon opening, low temperature sustains repression of embryonic-related genes in *prc2* mutants by preventing H2A.Z reduction. Additionally, our data suggests that H2A.Z contributes to gene repression through monoubiquitination. TARGET OF EARLY ACTIVATION TAGGED (EAT) 1 (TOE1) functions as a mediator of differential H2A.Z dynamics under varying temperatures. Our findings illuminate H2A.Zub’s compensatory role in the absence of H3K27me3 at low temperature, indicating cooperation between H2A.Z and H3K27me3 for *Arabidopsis* germination.

## Results

### Low ambient temperature restores the developmental defects of *prc2*

PRC2 loss-of-function triggers Polycomb callus formation during post-germination development (Figure S1A).^16^ Core PRC2 component mutations, such as *clf-28 swn-7*^36^ and *cdka;1-fertilization-independent endosperm (cdka;1-fie)*^17^, results in ‘callus’ formation in most plants (92.4% and 84.6%, respectively) at the optimal temperature (22°C). Interestingly, under low ambient temperature (16°C), 96.1% of *clf-28 swn-7* and 90.6% of *cdka;1-fie* seedlings develop into relatively regular plants, referred to as ‘plant,’ with rosette leaves, cauline leaves, and later infertile flower structures, despite their small and pale appearance (Figures 1A and 1B). To confirm these morphological differences, we conducted RNA-seq analysis using ‘callus’ and ‘plant’ samples from *clf-28 swn-7* seedlings grown at different temperatures. A significant number of genes, with 3,933 out of 20,261 expressed genes, exhibited differential expression between ‘callus’ and ‘plant’ samples (Figure 1C). In the ‘callus’ samples, genes related to somatic embryogenesis, lipid storage, seed maturation, and response to auxin were highly expressed (Figure 1D), including *ABI3*, *LEC1*, *CUP-SHAPED COTYLEDON1* (*CUC1*) and *LEC2* (Figure 1E). Conversely, ‘plant’ samples showed high expression of genes involved in chlorophyll biosynthesis, photosynthesis, and chloroplast thylakoid functions (Figure 1D), such as *PSI-N* (*PSAN*), *PSI-G* (*PSAG*), *RIBULOSE BISPHOSPHATE CARBOXYLASE SMALL CHAIN 1A* (*RBCS1A*) and *RUBISCO SMALL SUBUNIT 3B* (*RBCS3B*) (Figure 1F). The transcriptome aligns with morphological characteristics, providing further validation that low temperature effectively restores the developmental defects of *prc2* mutants.

**Figure 1.**
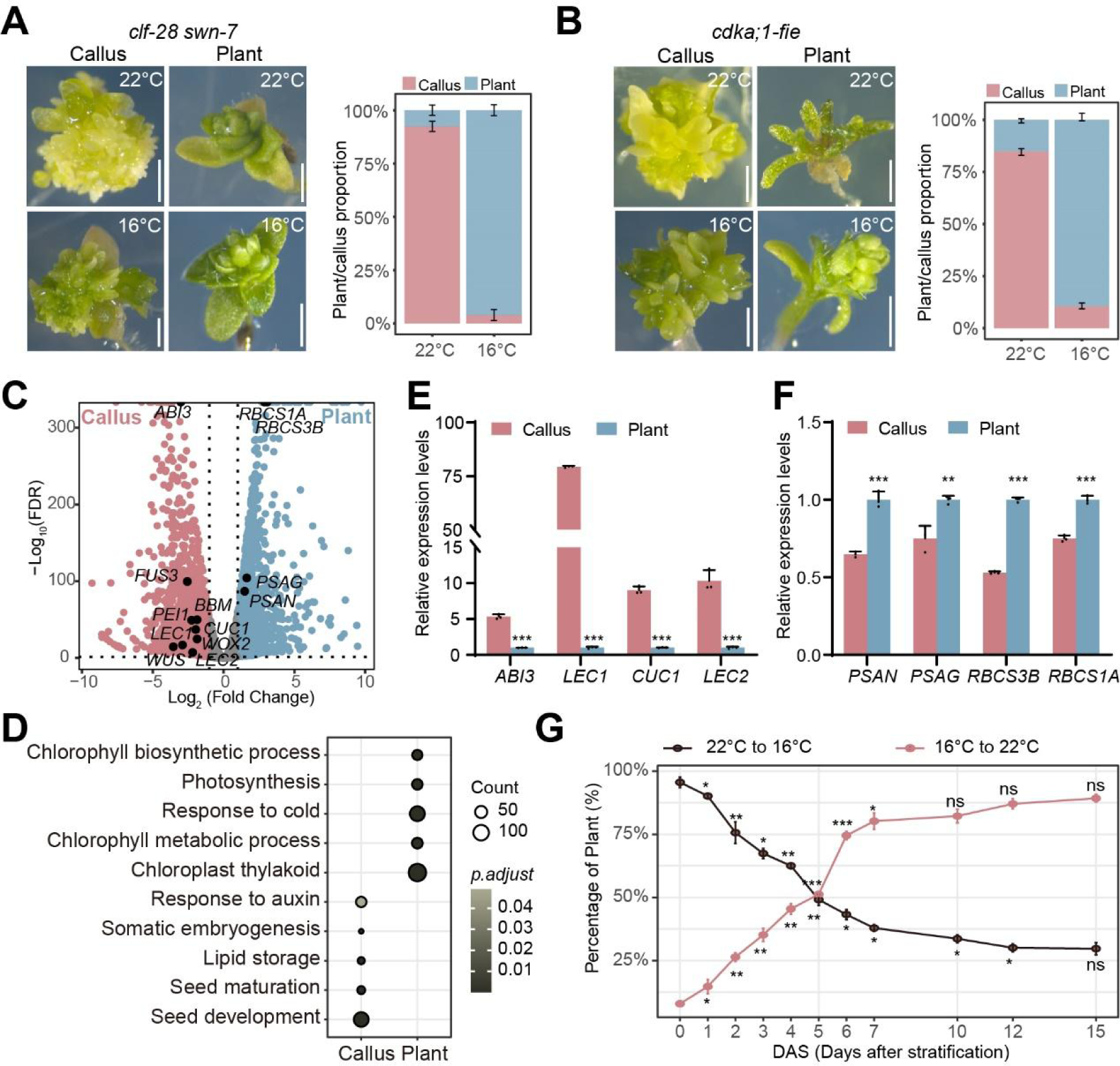
Low ambient temperature restored the developmental defects of *prc2*. (A and B) The ‘callus’ and ‘plant’ morphology and proportion of *clf-28 swn-7* (A) and *cdka;1-fie* (B) under different ambient temperatures, pictures were taken for 40 days grown at 22°C or 16°C. Scale bars, 1 mm. The accumulative histogram showed the proportion of ‘callus’ and ‘plant’ of *prc2* mutants at 22°C and 16°C. Data was mean ± SD from three biological replicates, n = 50 for each replicate. (C) Volcano plot of genes with elevated expression in ‘plant’ (blue) or ‘callus’ (pink) in *clf-28 swn-7* under different temperature. Gray dots, no difference between two samples. The known genes were labelled and shown in black dots. The cutoff of significance: Log_2_ (Fold change) > 1 or < −1; FDR < 0.05. (D) GO enrichment analysis of genes with elevated expression in ‘callus’ or ‘plant’ in *clf-28 swn-7*. (E and F) Relative expression levels of known marker genes in ‘callus’ (E) and ‘plant’ (F) as determined by qRT-PCR, expression in ‘plant’ was set as 1. Shown was mean ± SD of three biological replicates. Significance was calculated by Student’s *t*-test, ** *P* < 0.01; *** *P* < 0.001. (G) Temperature shift assay for *clf-28 swn-7*. The ‘plant’/’callus’ proportion of *clf-28 swn-7* was counted at 35 DAS. Data was mean ± SD from three biological replicates, n >30 for each replicate. Each time point was compared with the previous time point. Significance was calculated by Student’s *t*-test, * *P* < 0.05; ** *P* < 0.01; *** *P* < 0.001; ns, not significant (*P* > 0.05).

To define the developmental window influenced by ambient temperatures in shaping cell fate, we conducted a temperature shift assay. After stratification at 4°C, *clf-28 swn-7* seeds were initially grown at 22°C for varying days and then transferred to 16°C, or vice versa. The percentage of ‘plant’ morphology decreased with prolonged initial growth at 22°C and increased with prolonged growth at 16°C within the first 15 days after stratification (DAS) (Figure 1G). Notably, ‘callus’ and ‘plant’ percentages were similar, around 50% each, after the initial 5 DAS at either 22°C or 16°C. Subsequent to 7 DAS of initial growth at 16°C and 15 DAS of initial growth at 22°C, the changes in ‘callus’ and ‘plant’ percentages stabilized, suggesting that cell fate is primarily determined within the first 15 DAS (Figure 1G).

Considering multiple genes encoding the histone methyltransferase subunit of the PRC2 complex,^6^ we examined the genome-wide level of H3K27me3 in *clf-28 swn-7* seedlings under different ambient temperatures. ChIP-seq analysis confirmed a sharp decline in genome-wide H3K27me3 levels in *clf-28 swn-7* seedlings at both temperatures (Figures S1B and S1C). The finding suggests that the restoration of developmental defects in *clf-28 swn-7* at low temperature occurs independently of H3K27me3 recovery.

### Low ambient temperature re-silences ectopically expressed embryonic genes in *clf-28 swn-7*

To investigate morphological variations in *clf-28 swn-7* at different temperatures, we monitored post-germination development until distinct defects appeared. Our analysis, spanning five stages (S1-S5), revealed notable morphological diversification emerging at S3 with true leaf appearance, and pronounced differences at S4, marking the transition to callus or plant development (Figure 2A). Notably, the time of discernible phenotypic differences (S4) coincided with the stabilization of the ‘plant/callus’ ratio in the temperature shift assay, indicating that *clf-28 swn-7* cell fate determination occurs within 15 DAS.

**Figure 2.**
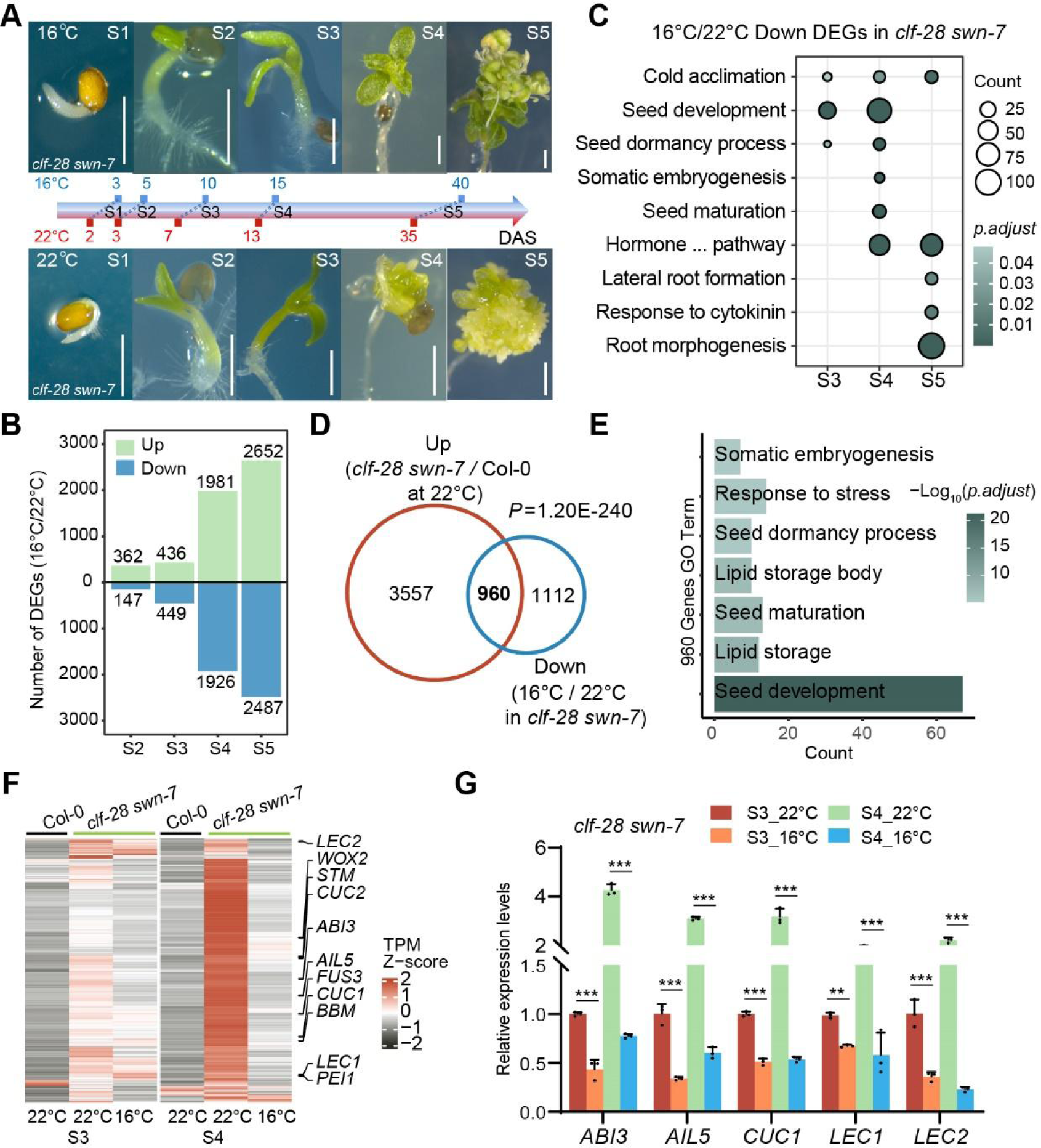
The transcriptome analysis of *clf-28 swn-7* under different temperature at various developmental stages. (A) The morphological diverse of *clf-28 swn-7* at five different developmental stages under 22°C and 16°C. Growth days corresponding to different developmental stages are written in blue (16°C) and red (22°C), respectively. DAS: days after stratification. (B) The number of down- and up-regulated genes in *clf-28 swn-7* at 16°C compared with 22°C from stage S2 to S5 stages. (C) GO enrichment analysis of down-regulated genes in *clf-28 swn-7* at 16°C compared to 22°C from stage S3 to S5. (D) Venn diagram showing 960 genes with elevated expression in *clf-28 swn-7* as compared to Col-0 at 22°C but down-regulated at 16°C when compared to 22°C. (E) GO enrichment analysis of the 960 genes from (D). (F) Heatmap showing the expression pattern of the 960 genes from (D) in Col-0 and *clf-28 swn-7* under different temperatures at S3 and S4 stages, with Z-score of normalized TPM determined by RNA-seq. (G) Relative expression levels of *ABI3*, *AIL5*, *CUC1*, *LEC1* and *LEC2* at S3 and S4 stages in *clf-28 swn-7* under different temperatures, as determined by qRT-PCR. Expression in *clf-28 swn-7* under 22°C at S3 stage was set as 1. Shown was mean ± SD of three biological replicates. Significance was calculated by Student’s *t*-test, ** *P* < 0.01; *** *P* < 0.001.

To delve into the transcriptome dynamics underlying cell fate changes at different temperatures, RNA-seq data for *clf-28 swn-7* from S2-S5 stages was analyzed. The number of differentially expressed genes (DEGs) between temperatures sharply increased from S3 to S5 in *clf-28 swn-7* (Figure 2B), consistent with morphological divergence. PCA analysis revealed progressive transcriptional differences between temperatures, becoming pronounced at S3 (Figure S2A). GO enrichment analysis highlighted down-regulated genes at 16°C involved in seed development, somatic embryogenesis, and lateral root development from S3 to S5 (Figure 2C), corresponding to callus formation process.^37–39^ Whereas, genes up-regulated at 16°C are involved in cell wall biogenesis, chloroplast development and meiotic cell cycle from S3 to S5 (Figure S2B), corresponding to plant development. Moreover, at equivalent developmental stages (S3 and S4) in Col-0, minimal transcriptional differences were observed between temperatures, in stark contrast to *clf-28 swn-7* (Figure S2C). Differentially expressed genes between two temperatures at S3 and S4 in Col-0 were mostly enriched in secondary metabolism processes (Figure S2D). These results underscore PRC2 absence accentuates low ambient temperature effects on cell fate. Notably, PCA analysis showed that transcriptome of *clf-28 swn-7* at 16°C was more similar to Col-0 than at 22°C at S4 (Figure S2C), aligning with the obvious morphological recovery of *clf-28 swn-7* under 16°C at S4 stage (Figure 2A).

At S3 and S4 stages, 960 genes were up-regulated in *clf-28 swn-7* compared to Col-0 at 22°C but re-silenced at 16°C in *clf-28 swn-7* (Figure 2D). These genes, enriched in seed development and somatic embryogenesis (Figure 2E), including *ABI3*, *AINTEGUMENTA-LIKE 5* (*AIL5*), *CUC1*, *LEC1* and *LEC2* (Figures 2F and 2G), are known to promote polycomb callus formation when ectopically expressed.^40–42^ The re-silencing of these cell fate-related genes at S3 and S4 under low ambient temperature likely inhibits the callus formation in *clf-28 swn-7*.

### Low ambient temperature reduced PRC2 binding at target genes

To understand the diminished expression of cell fate determining genes in *clf-28 swn-7* at 16°C, we investigated PRC2 binding in Col-0 under normal and low temperatures, given that these genes are regulated by PRC2. FIE as the unique core subunit of PRC2,^17^ was chosen to represent PRC2 binding by ChIP-seq. Interestingly, the number of FIE binding targets and peaks decreased greatly at 16°C compared to 22°C, with a notable weakening in binding strength (Figures 3A - C and S3A). This change doesn’t seem linked to altered FIE expression, although the level of the PRC2 components *EMBRYONIC FLOWER 2* (*EMF2*) and *ARABIDOPSIS MULTICOPY SUPRESSOR OF IRA1* (*MSI1*) expression slightly increased, as shown by RNA-seq data (Figure S3B). 3,970 genes lost FIE binding at 16°C compared to 22°C in Col-0 (Figure 3C). Unexpectedly, only 51 of these 3,970 (∼1.3%) genes (group I) showed increased expression and decreased H3K27me3 level (Figures 3D and S3C). Surprisingly, in the gene sets where FIE-binding abolished but did not cause up-regulation (group II+III, n=3,919), H3K27me3 levels also slightly decreased around TSS regions at 16°C compared to 22°C (Figure 3E). This led us to postulate existence of compensatory chromatin modifications that maintain normal *Arabidopsis* development when PRC2 or H3K27me3 levels declined.

**Figure 3.**
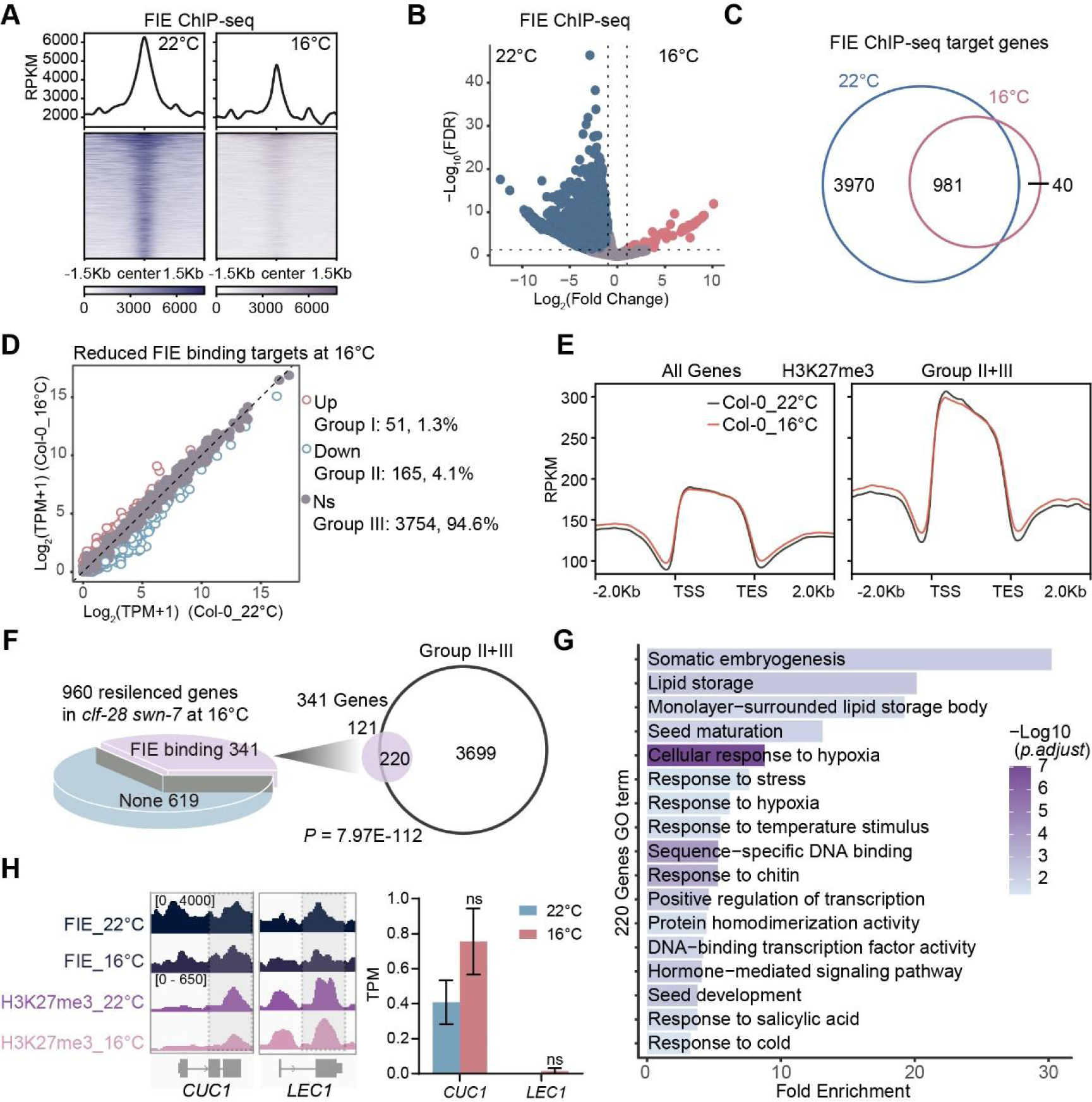
ChIP-seq analysis to test chromatin occupancy of FIE at 22°C and 16°C in Col-0. (A) Profiles and heatmaps showing peak enrichment for PRC2 (FIE) binding dynamic at 22°C and 16°C in Col-0. (B) Volcano plot of FIE binding peaks at different temperatures, with colored dots indicating elevated binding at either 22°C (blue) or 16°C (pink). Gray dots, no difference. The cutoff of significance was Log_2_ (Fold change) > 1 or < −1; FDR < 0.05. (C) Venn diagram showing number of FIE target genes at 22°C and 16°C in Col-0. (D) The dot plot showing Varied expression pattern of 3970 abolished FIE-binding targets from (C) at 16°C and 22°C in Col-0. Group I: up regulated at 16°C /22°C (pink); Group II: down regulated (blue); Group III: no significant change (gray). (E) ChIP-seq profile of H3K27me3 along genic region of all genes or abolished FIE-binding but not up-regulated genes (Group II + III, panel D) at 22°C and 16°C in Col-0. TSS, transcription start site; TES, transcription end site. (F) Pie chart (left) showing number of FIE targets in the 960 genes from (Figure 2D). And venn diagram (right) showing the overlap between the 341 FIE targets and non-up regulated genes (Group II+III) from (D). Fisher exact-test, *P* =7.97E-112. (G) GO enrichment analysis of the 220 genes from (F). (H) IGV browser view of ChIP-seq signals of FIE and H3K27me3 (left) at *CUC1* and *LEC1* loci under 22°C and 16°C. Bar graph showing the expression pattern (right) of *CUC1* and *LEC1* at 22°C and 16°C. Data was mean ± SD of 3 biological replicates from RNA-seq. Student’s *t*-test, ns, no significant changes.

Among the 960 re-silenced genes in *clf-28 swn-7* at 16°C compared to 22°C (Figure 2D), 341 were direct targets of FIE binding according to ChIP-seq analysis (Figure 3F). Notably, these 341 genes showed a significant correlation with the abolished FIE-binding but not up-regulated genes (group II+III) in Col-0 (Fisher exact-test; *P* = 7.97E-112), with 220 overlapped genes (Figure 3F). These genes are enriched in seed development and somatic embryogenesis, including known genes contributing to callus information (Figure 3G), like *LEC1* and *CUC1*.^43,44^ Representative genes *LEC1* and *CUC1* exhibited attenuated FIE binding and reduced H3K27me3 accumulation, but no significant change of expression levels at 16°C compared to 22°C in Col-0 (Figure 3H). Besides, GO enrichment analysis indicated not up-regulated genes (group II+III) in Col-0 were also enriched in processes like seed development, root morphogenesis, and respond to auxin (Figure S3D). These results suggest that for the 341 genes in *clf-28 swn-7* at 16°C, the mechanism likely resembles that in Col-0—where silencing of PRC2 targets occurs through other chromatin modifications in the absence of PRC2.

### PRC1 and H2A.Z collaborate in restoring growth defects of *clf-28 swn-7* at low ambient temperature

During germination, key genes involved in seed development and maturation including master regulators *ABI3*, *FUS3*, *LEC2*, *LEC1*,^43^ as part of the previously identified 341 genes, are sequentially repressed by PRC1 and PRC2.^14^ To explore whether PRC1 compensates for PRC2 absence at 16°C, we generated a *ring1a clf-28 swn-7* triple mutant by crossing *clf-28 swn-7* with *ring1a*, a PRC1 subunit mutant.^44^ The developmental defects in *ring1a clf-28 swn-7* were more severe, with only 51.3% displaying ‘plant’ morphology at 16°C compared to *clf-28 swn-7* (Figures 4A and S4A), indicating the requirement of PRC1 for restoring *clf-28 swn-7* growth defects under low ambient temperature.

**Figure 4.**
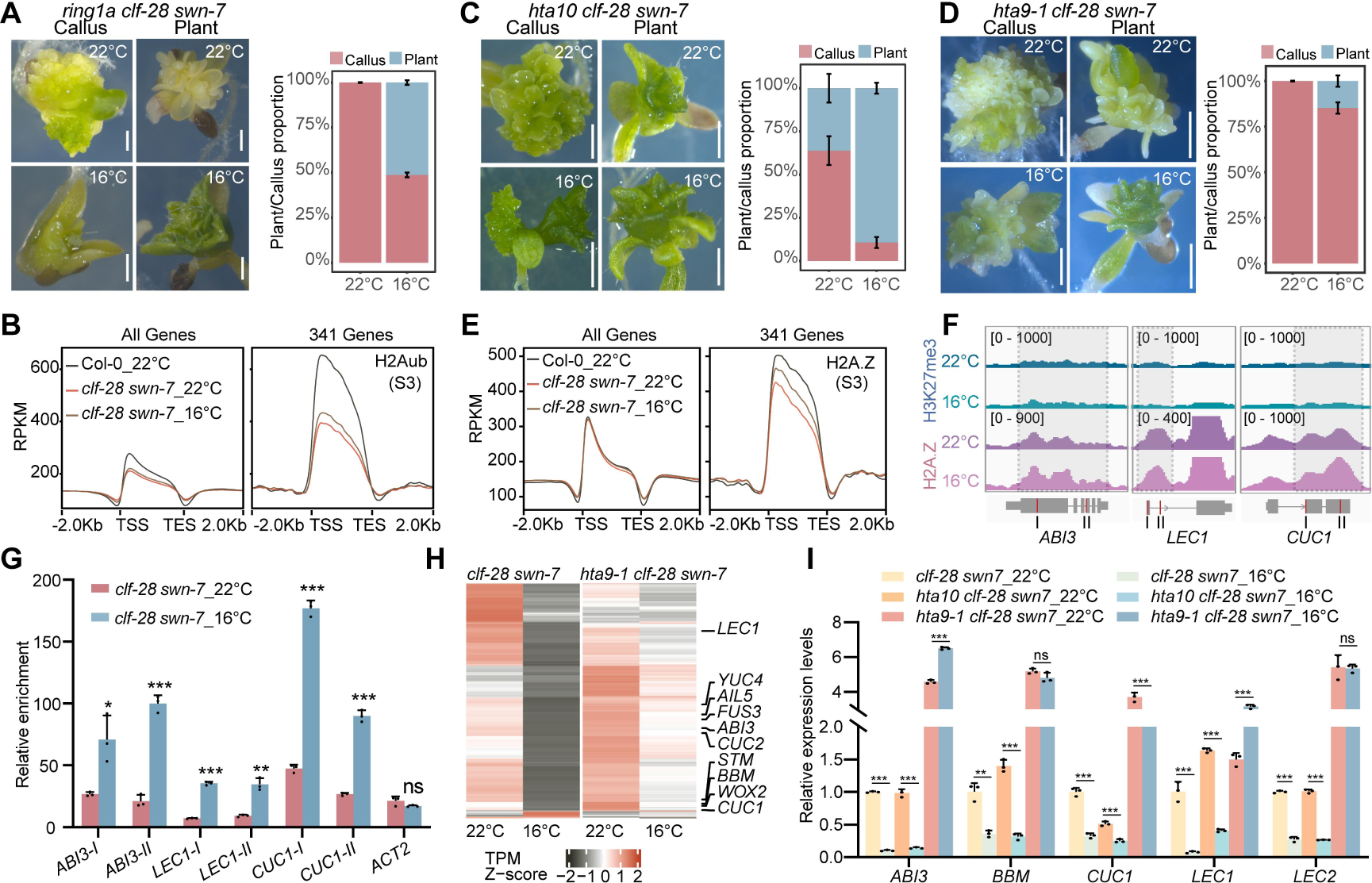
PRC1 and H2A.Z but not H2A are involved in the restoration of *clf-28 swn-7* growth defect at low temperature. (A) The ‘callus’ and ‘plant’ morphology and proportion of *ring1a clf-28 swn-7* under different ambient temperatures, pictures were taken for 40 days grown at 22°C or 16°C. Scale bars, 1 mm. The accumulative histogram showed the proportion of ‘callus’ and ‘plant’ at 22°C and 16°C. Data was mean ± SD from three biological replicates, n >30 for each replicate. (B) ChIP-seq profile of H2Aub along genic region of all genes and the 341 genes (Figure 3F) in *clf-28 swn-7* at S3 stage under 22°C and 16°C, as well as in Col-0 at 22°C. TSS, transcription start site; TES, transcription end site. (C and D) The ‘callus’ and ‘plant’ morphology and proportion of *hta10 clf-28 swn-7* (C) and *hta9-1 clf-28 swn-7* (D) under different ambient temperatures, pictures were taken for 40 days grown at 22°C or 16°C. Scale bars, 1 mm. The accumulative histogram showed the proportion of ‘callus’ and ‘plant’ at 22°C and 16°C. Data was mean ± SD from three biological replicates, n >30 for each replicate. (E) ChIP-seq profile of H2A.Z along genic region of all genes and the 341 genes (Figure 3F) in *clf-28 swn-7* at S3 stage under 22°C and 16°C, as well as in Col-0 at 22°C. TSS, transcription start site; TES, transcription end site. (F) IGV browser view of H2A.Z and H3K27me3 signal at *ABI3, LEC1* and *CUC1* at 22°C and 16°C in *clf-28 swn-7*. (G) Enrichment of H2A.Z at *ABI3*, *LEC1* and *CUC1* under 22°C and 16°C in *clf-28 swn-7*, as measured by CUT&Tag-qPCR. *ACT2* as a negative control. Red lines in schematic gene structures in (F) indicate regions examined by CUT&Tag-qPCR. Data was mean ± SD of three replicates. Significance was calculated by Student’s *t*-test, * *P* < 0.05; ** *P* < 0.01; *** *P* < 0.001; ns, no significance. (H) Heatmap showing transcription dynamic of the 341 genes (Figure 3F) under 22°C and 16°C at S4 stage in *clf-28 swn-7* and *hta9-1 clf-28 swn-7*, with Z-score of normalized average TPM of three replicates determined by RNA-seq. The known cell fate regulator genes were listed. (I) Relative expression level of *ABI3*, *BBM*, *CUC1*, *LEC1* and *LEC2* at S4 stage in *clf-28 swn-7*, *hta10 clf-28 swn-7*, and *hta9-1 clf-28 swn-7* under different temperatures, as determined by qRT-PCR. Expression in *clf-28 swn-7* under 22°C at S4 stage was set as 1. Shown was mean ± SD of three biological replicates. Significance was calculated by Student’s *t*-test, ** *P* < 0.01; *** *P* < 0.001; ns, no significance.

Research generally posits that PRC1 suppresses gene expression through H2A monoubiquitination.^8,45,46^ ChIP-seq analysis showed a significant reduction in H2Aub level at 22°C and 16°C in *clf-28 swn-7*, with Col-0 at 22°C as a control. Interestingly, there was a slight increase in H2Aub in *clf-28 swn-7* at 16°C compared to 22°C, both across the genome and specifically at the 341 genes (Figure 4B). To investigate if this non-specific H2Aub increase contributes to the re-silencing of the 341 genes at 16°C, we generated a *hta10 clf-28 swn-7* triple mutant by crossing *clf-28 swn-7* with *histone h2a 10* (*hta10*), a mutation in one H2A coding gene. Although H2Aub level reduced in *hta10* mutants compared to Col-0 (Figure S4B), the *hta10 clf-28 swn-7* triple mutant exhibited similar (*P* = 0.0452) ‘plant/callus’ proportion (89.3% of plant, 10.7% callus) to *clf-28 swn-7* (96.1% of plant, 3.9% callus) at 16°C, and even higher ‘plant’ ratios of *hta10 clf-28 swn-7* (36.1%) than *clf-28 swn-7* (7.6%) at 22°C (Figures 4C and S4A), suggesting that PRC1 may not function through H2Aub in this context.

It has been reported that PRC1 can also catalyze the monoubiquitination of H2A.Z to mediate transcriptional repression.^9^ Accordingly, we generated *hta9-1 clf-28 swn-7* triple mutant by crossing *clf-28 swn-7* with *histone h2a protein 9-1* (*hta9-1*), a mutation affecting one of H2A.Z coding genes.^45^ Remarkably, only about 14.8% of *hta9-1 clf-28 swn-7* developed into ‘plant’ at 16°C, significantly lower than *clf-28 swn-7* (Figures 4D and S4A). Global profiling of H2A.Z at two temperatures in *clf-28 swn-7*, using Col-0 at 22°C as a control, revealed a specific increase in H2A.Z levels on the 341 genes at 16°C compared to 22°C in *clf-28 swn-7* (Figure 4E). This increase wasn’t observed genome-wide (Figure 4E). *ABI3, LEC1 and CUC1* as representative cases showed increase of H2A.Z in *clf-28 swn-7* at 16°C compared to 22°C, without H3K27me3 restoration (Figure 4F). CUT&Tag-qPCR^47^ further validated the pattern on these genes (Figures 4G). Moreover, RNA-seq analysis revealed the transcription levels of the 341 genes were higher in *hta9-1 clf-28 swn-7* than *clf-28 swn-7* at 16°C at the S4 stage (Figure 4H). The expression of key genes like *ABI3*, *BBM*, *CUC1*, *LEC1* and *LEC2* was significant higher in *hta9-1 clf-28 swn-7* than in *clf-28 swn-7* or *hta10 clf-28 swn-7* (Figure 4I) at 22°C, as assayed by RT-qPCR. Importantly, at 16°C, these genes remained active in *hta9-1 clf-28 swn-7*, different from *clf-28 swn-7* or *hta10 clf-28 swn-7*, where they were silenced (Figures 4I and S4C). Besides, GO enrichment analysis highlighted up-regulated genes of *hta9-1 clf-28 swn-7* compared to *clf-28 swn-7* at 16°C involved in somatic embryogenesis, seed maturation and dormancy, and seed development (Figure S4D), corresponding to the embryonic traits. This underscores the role of H2A.Z in repressing embryonic-related genes and highlights its importance in low temperature-induced silencing of these genes and restoration of cell fate. Thus, PRC1’s role likely relies on H2A.Z rather than H2A in this context.

### PRC1-catalyzed H2A.Zub and PRC2-mediated H3K27me3 coordinately silencing seed development program during post-germination

It is notable that the morphological characteristics of *hta9-1 clf-28 swn-7* before cotyledon expansion resemble those of the robust PRC1 mutant *atbmi1a/b/c* described previously,^14^ showing white cotyledons and swollen root (Figure 5A). The DEGs in *hta9-1 clf-28 swn-7* compared to *clf-28 swn-7* are significantly correlated with the DEGs of *atbmi1a/b/c* compared to Col-0 at 10 DAG (Figure S5A),^13^ suggesting potential cooperation between H2A.Z and PRC1 in early seedling development post-germination.

**Figure 5.**
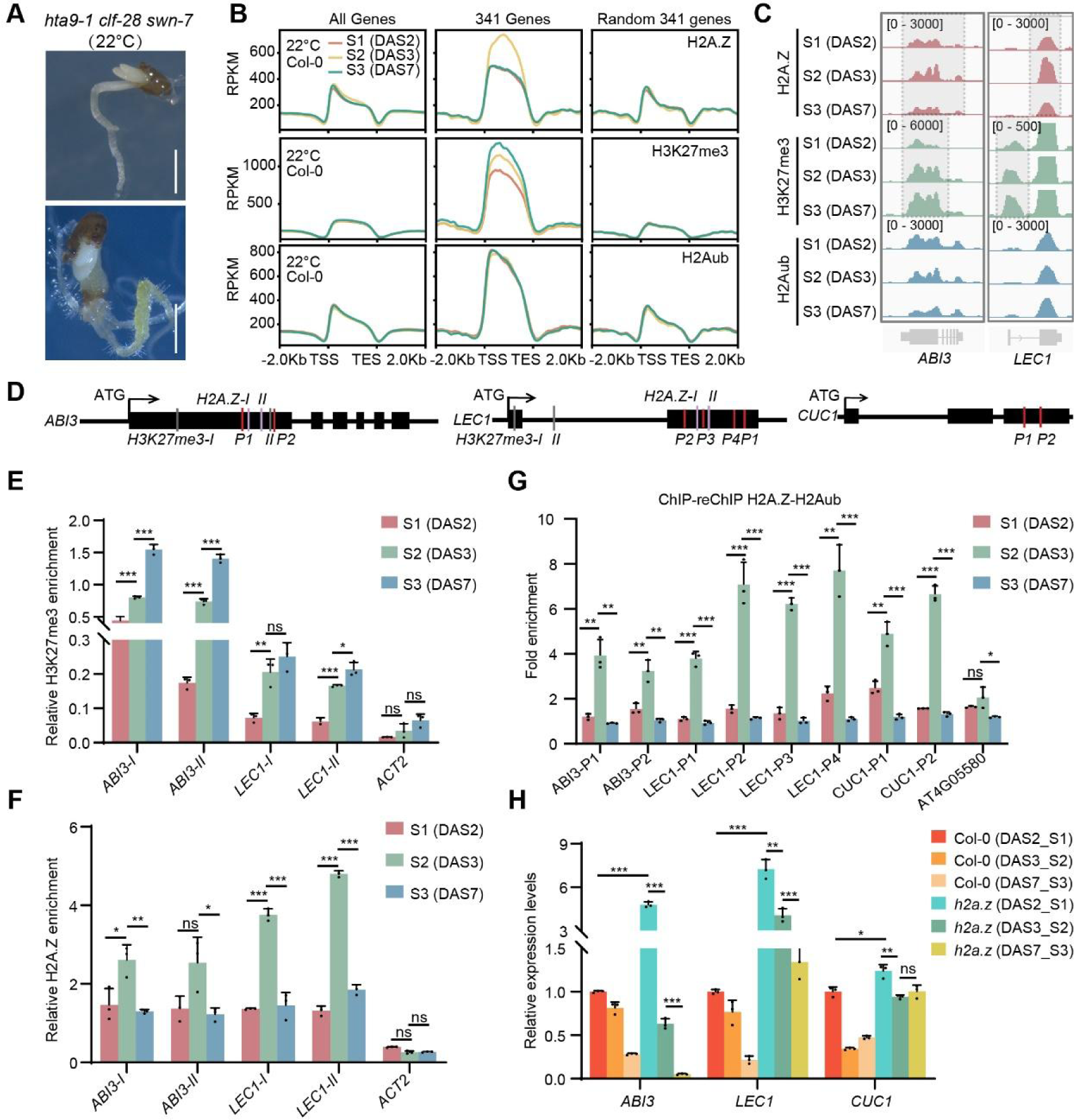
PRC1-catalyzed H2A.Zub and PRC2-mediated H3K27me3 coordinately inhibit seed development program post-germination. (A) The *hta9-1 clf-28 swn-7* showed abnormal embryonic cotyledons at 22°C. Scale bars, 1 mm. (B) ChIP-seq profile of H2A.Z, H3K27me3 and H2Aub along genic region of all genes, the 341 genes (Figure 3F) and the other random 341 genes from 2 DAS (S1) to 7 DAS (S3) in Col-0 at 22°C. TSS, transcription start site; TES, transcription end site. (C) IGV browser view of H2A.Z, H3K27me3 and H2Aub signal at *ABI3* and *LEC1* at three time points in Col-0 at 22°C. (D) Schematic structures of *ABI3*, *LEC1*, and *CUC*1. Arrows indicate transcription start sites. Lines indicate regions examined by H3K27me3 (gray) ChIP-qPCR, H2A.Z (pink) ChIP-qPCR and re-ChIP-qPCR (red). (E and F) ChIP-qPCR assays showing enrichment of H3K27me3 (E) and H2A.Z (F) at *ABI3* and *LEC1* from 2 DAS (S1) to 7 DAS (S3) in Col-0 at 22°C. *ACT2* as a negative control. Data was mean ± SD of three replicates. Significance was calculated by Student’s *t*-test, * *P* < 0.05; ** *P* < 0.01; *** *P* < 0.001; ns, no significance. (G) ReChIP-qPCR assay showing enrichment of H2A.Zub at *ABI3*, *LEC1* and *CUC1* from 2 DAS (S1) to 7 DAS (S3) in Col-0 at 22°C. *AT4G05580* as a negative control. Data was mean ± SD of three replicates. Significance was calculated by Student’s *t*-test, * *P* < 0.05; ** *P* < 0.01; *** *P* < 0.001; ns, no significance. (H) Relative expression level of A*BI3*, *LEC1* and *CUC1* from 2 DAS (S1) to 7 DAS (S3) in Col-0 and *h2a.z* mutant, as determined by qRT-PCR. Expression in Col-0 at 2 DAS (S1) was set as 1. Shown was mean ± SD of three biological replicates. Significance was calculated by Student’s *t*-test, * *P* < 0.05; ** *P* < 0.01; *** *P* < 0.001; ns, no significance.

In Col-0, seed germination occurs at 2 DAS, cotyledons open at 3 DAS, and true leaves emerge at 7 DAS. The three time points correspond to S1/S2/S3 developmental stages of *clf-28 swn-7* mutant, respectively (Figure 2A). In Col-0 at 22°C, ChIP-seq analysis revealed H2A.Z specifically increased from S1 to S2 (2 DAS to 3 DAS) and then declined at S3 (7 DAS) on the 341 genes, while H3K27me3 gradually increased from S1 to S3 (2 DAS to 7 DAS) on the same genes (Figure 5B). *ABI3* and *LEC1* as representative cases showed the pattern of H2A.Z and H3K27me3 from S1 to S3 (2 DAS to 7 DAS) (Figure 5C). ChIP-qPCR results confirmed these changes at *ABI3* and *LEC1* loci (Figures 5D-F). However, H2Aub levels on the 341 genes showed no significant changes, suggesting it may not be involved in inhibiting seed development post-germination (Figure 5B). At a genome-wide level, all three histone modification marks did not exhibit dramatic changes from S1 to S3 (2 DAS to 7 DAS) (Figure 5B). These dynamic patterns indicate that both H2A.Z and H3K27me3 play coordinated roles in silencing the seed development program, albeit likely with distinct functions.

We then investigated whether PRC1 catalyzes H2A.Zub to inhibit seed maturation program post-germination. Due to the lack of a specific antibody for H2A.Zub, we employed the H2Aub antibody (Millipore, 05-678), which recognizes both H2Aub and H2A.Zub.^46^ H2A.Z antibody was first used to enrich H2A.Z-containing nucleosomes, followed by re-ChIP with the H2Aub antibody to enrich mono-ubiquitinated H2A.Z (Figure S5B, see method for details). ChIP-re-ChIP confirmed enrichment of H2A.Zub at *ABI3*, *LEC1* and *CUC1* loci, with levels increasing from S1 to S2 (2 DAS to 3 DAS) and then decreasing from S2 to S3 (3 DAS to 7 DAS) (Figure 5G), consistent with the H2A.Z pattern (Figure S5C). Furthermore, expression of *ABI3*, *LEC1* and *CUC1* exhibited a gradual reduction in Col-0 (Figure 5H), aligning with the repression of seed maturation program post-germination.^18,47^ But in *h2a.z* (*hta8-1 hta9-1 hta11-1*) mutant,^29^ these genes expressed higher compared to Col-0 at S1 (2 DAS) (Figure 5H), indicating H2A.Zub’s involvement in gene repression post-germination. However, even without H2A.Z, the repression of these genes from S1 to S3 (2 DAS to 7 DAS) still occurred (Figure 5H), suggesting a potential contribution of H3K27me3 to repression of these genes. Indeed, the expression of *ABI3*, *LEC1* and *CUC1* was activated from S2 to S3 (3 DAS to 7 DAS) stages in *clf-28 swn-7* (Figure S5D).

The results indicate that during germination process, PRC1-mediated H2A.Zub initially predominantly represses key embryonic genes. While H2A.Zub decreases, the gradual increase of PRC2-mediated H3K27me3 compensates for this reduction, ensuring continuous suppression of embryonic genes, thus maintaining normal post-germination development.

### Low temperature slowdown H2A.Z turnover specifically at cell fate related genes via TOE1

To delve deeper into why H2A.Z specifically accumulates at the 341 cell fate-related genes in *clf-28 swn-7* under low temperature (Figure 4E), we examined H2A.Z dynamics at S2, S2.5 (5 DAS at 22°C and 6 DAS at 16°C) and S3 time points in *clf-28 swn-7* by ChIP-seq. At 22°C, H2A.Z peaked at S2, declined sharply at S2.5, and gradually decreased at S3 specifically on the 341 genes (Figure 6A). In contrast, at 16°C, H2A.Z levels remained steady, sustaining a high level from S2 to S3 (Figure 6A), indicating a potential slowdown in H2A.Z turnover in *clf-28 swn-7* under low temperature. At the genome-wide level, no significant change in H2A.Z occupancy was observed from S2 to S3 at either 22°C or 16°C (Figure 6A).

**Figure 6.**
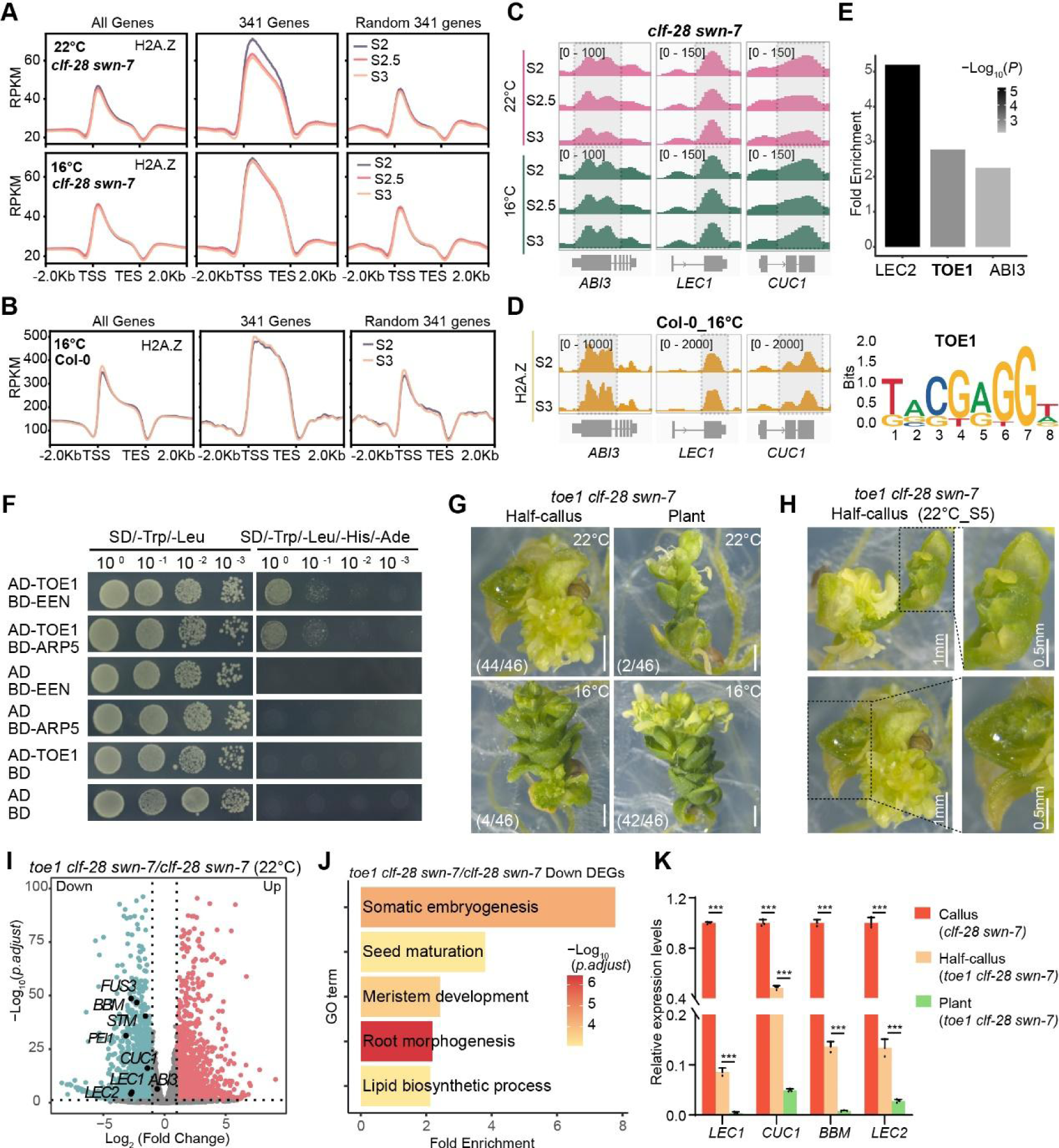
Low temperature specifically slowdown H2A.Z turnover at key targets. (A) ChIP-seq profile of H2A.Z along genic region of all genes, the 341 genes (Figure 3F) and the other random 341 genes from S2 to S3 at 22°C and 16°C in *clf-28 swn-7*. TSS, transcription start site; TES, transcription end site. (B) ChIP-seq profile of H2A.Z along genic region of all genes, the 341 genes (Figure 3F) and the other random 341 genes from S2 to S3 at 16°C in Col-0. (C) IGV browser view of H2A.Z signal at *ABI3*, *LEC1* and *CUC1* from S2 to S3 at 22°C and 16°C in *clf-28 swn-7*. (D) IGV browser view of H2A.Z signal at *ABI3*, *LEC1* and *CUC1* from S2 to S3 at 16°C in Col-0. (E) Barplot showing enrichment of top three transcription factors from promoter region of the 341 genes. Motif sequence of TOE1 was shown. (F) Y2H assay showing the interaction between TOE1 and INO80-C (EEN and ARP5). (G) The ‘half-callus’ and ‘plant’ morphology of *toe1 clf-28 swn-7* under different ambient temperatures, pictures were taken for 35 days grown at 22°C or 16°C. Scale bars, 1 mm. The proportion of ‘half-callus’ and ‘plant’ at 22°C and 16°C was shown. (H) ‘Half-callus’ morphology of *toe1 clf-28 swn-7*, pictures were taken for 35 days at 22°C. Scale bar is as indicated. (I) Volcano plot of genes with lower (blue) or higher (pink) expression in *toe1 clf-28 swn-7* compared to *clf-28 swn-7* at 22°C. Gray dots, no difference between two samples. The known genes were labelled and shown in black dots. The cutoff of significance: Log_2_ (Fold change) > 1 or < −1; *p.adjust* < 0.05. (J) GO enrichment analysis of the down-regulated genes in *toe1 clf-28 swn-7* compared to *clf-28 swn-7* at 22°C. (K) Relative expression levels of *LEC1*, *CUC1*, *BBM* and *LEC2* in ‘callus’ of *clf-28 swn-7*, ‘half-callus’ and ‘plant’ of *toe1 clf-28 swn-7*, as determined by qRT-PCR. Expression in ‘callus’ of *clf-28 swn-7* was set as 1. Shown was mean ± SD of three biological replicates. Significance was calculated by Student’s *t*-test, *** *P* < 0.001.

In Col-0 at 22°C, H2A.Z level also declined from S2 to S3 (Figure 5B). Interestingly, at 16°C, H2A.Z did not decrease from S2 (5 DAS at 16°C) to S3 (14 DAS at 16°C) at the cell fate-related 341 genes (Figures 6B). Quantitative analysis confirmed a significant decrease in H2A.Z levels from S2 to S3 at 22°C but not at 16°C for these genes (Figure S6A). The results indicated a temperature-regulated slowdown of H2A.Z turnover at these genes, irrespective of the presence of PRC2. By contrast, increase of H3K27me3 from S2 to S3 at the 341 genes was maintained at 16°C (Figure S6B) as 22°C (Figure 5B). Representative cases from 341 genes such as *ABI3*, *LEC1* and *CUC1* illustrate the dynamic changes of H2A.Z in *clf-28 swn-7* and Col-0 at two ambient temperatures (Figures 6C and 6D). Thus, low ambient temperature specifically slows down H2A.Z turnover at genes related to cell-fate determination.

We wonder how the low temperature specifically regulated H2A.Z dynamics at the 341 genes? Analysis of the chromatin accessible regions within the promoter regions of the H2A.Z direct targets of the 341 genes revealed the top three enriched transcription factor binding motifs: LEC2, TOE1, and ABI3 (Figure 6E, see method for details). Protein interaction assays including Y2H (Yeast two-hybrid) and BiFC (Bimolecular fluorescence complementation) showed that only TOE1 directly interacted with H2A.Z chaperone INO80-C (EIN6 ENHANCER (EEN) and ACTINRELATED PROTEIN 5 (ARP5)) (Figures 6F and S6C-E), which have been reported to participate in H2A.Z turnover.^30^ To further confirm the function of TOE1 in regulating H2A.Z dynamics specifically at those 341 genes, we generated *toe1 clf-28 swn-7* triple mutant by crossing *toe1 toe2* ^48^ with *clf-28 sw-7.* At 22°C, most triple mutants (95.7%, 44/46) exhibited “half-callus” characteristics with both leaf and callus structure in one plant (Figures 6G, 6H and S6F). RNA-seq analysis showed that key embryonic-related genes like *BBM*, *CUC1*, *LEC1* and *LEC2*, were significantly down-regulated in “half-callus” of *toe1 clf-28 swn-7* compared to “callus” of *clf-28 swn-7* at S5 stage and RT-qPCR confirmed the results (Figures 6I and 6K). GO enrichment analysis revealed down-regulated genes in *toe1 clf-28 swn-7* compared to *clf-28 swn-7* enriched in somatic embryogenesis and seed maturation (Figure 6J). Under 16°C, the majority of triple mutants (91.3%, 42/46) still developed into “plant”, with minimal expression of embryonic-related genes (Figures 6G and 6K). The results suggested that TOE1 is involved in mediating the restoration of *clf-28 swn-7* via influencing the transcription level of key embryonic genes.

### TOE1 responds to ambient temperature and regulates H2A.Z dynamics at embryonic genes

We further explored the regulation of TOE1 and H2A.Z dynamics specifically at embryonic genes and how TOE1 responds to ambient temperature to influence these dynamics. A ChIP-qPCR assay using *mTOE1-Flag* ^48^ confirmed the direct binding of TOE1 to the chromatin regions of *LEC1*, *LEC2* and *PEI1*, from the 341 genes, at both S2 (3 DAS) and S3 (7 DAS) stages at 22°C (Figures 7A and 7B). To investigate whether the interaction between TOE1 and INO80-C is associated with H2A.Z removal, we analyzed the H2A.Z dynamics in *toe1 toe2*^48^ double mutant. In this mutant, H2A.Z level didn’t reduce but were even higher from S2 to S3 stage at the 341 genes at 22°C (Figure 7C), unlike in Col-0. Representative cases, such as *LEC1*, *LEC2* and *PEI1* illustrate the maintained high level of H2A.Z in *toe1 toe2* compared to Col-0 from S2 to S3 stage at 22°C (Figure 7D). These results suggest a connection between TOE1 and eviction of H2A.Z at embryonic genes during post-germination stages.

**Figure 7.**
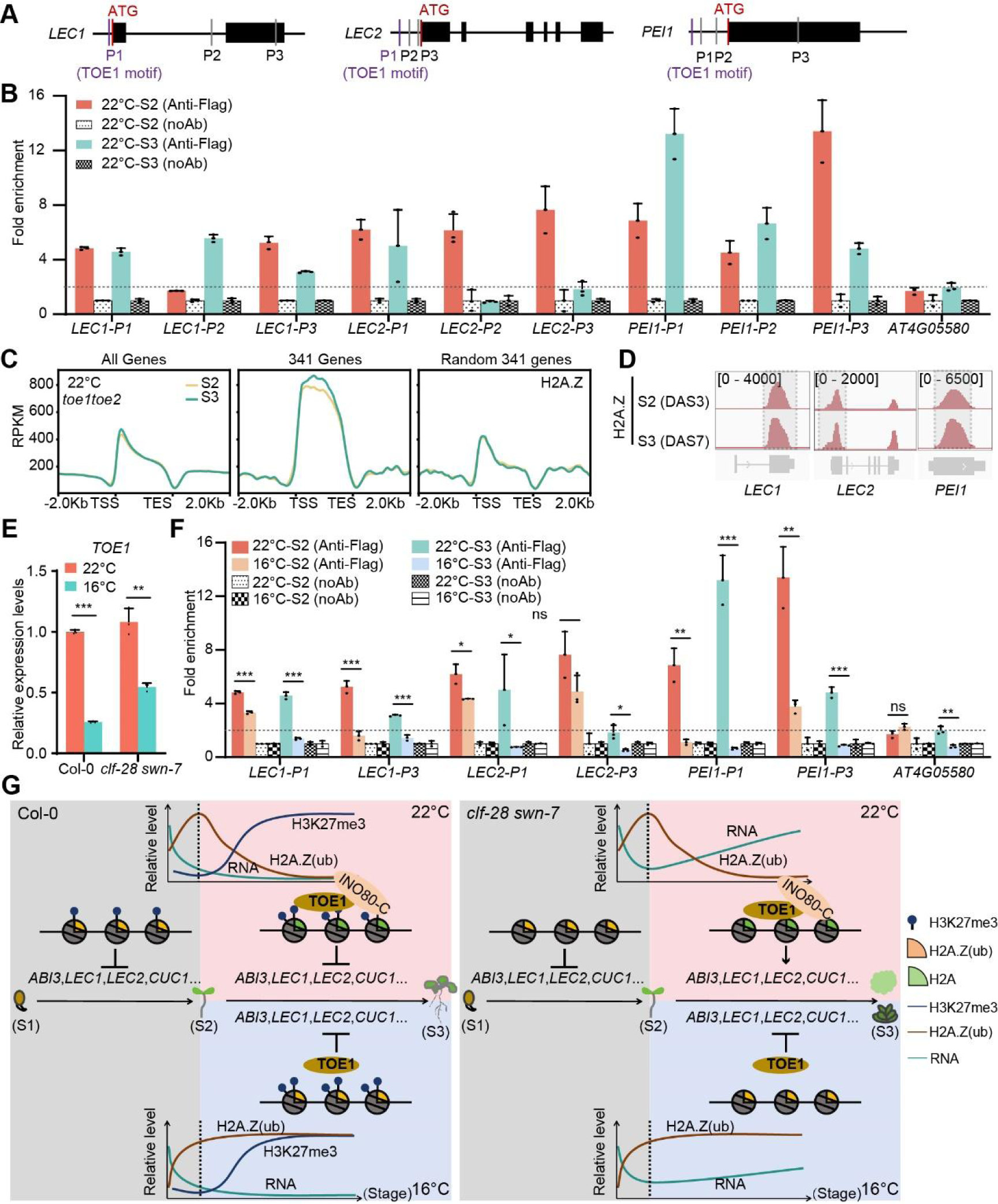
Temperature regulates H2A.Z dynamics to shape cell fate by TOE1. (A) The diagrams showing the locations of specific fragments that were amplified by ChIP-qPCR with specific primers in (B) and (F). The purple lines indicate the motif of TOE1. (B) ChIP-qPCR assay showing enrichment of TOE1 at *LEC1*, *LEC2* and *PEI1* at S2 and S3 stages at 22°C in Col-0. *AT4G05580* was a negative control. Data was mean ± SD of three biological replicates. (C) ChIP-seq profile of H2A.Z along genic region of all genes, the 341 genes (Figure 3F) and the other random 341 genes from S2 (3 DAS) to S3 (7 DAS) in *toe1 toe2* at 22°C. TSS, transcription start site; TES, transcription end site. (D) IGV browser view of H2A.Z signal at *LEC1*, *LEC2* and *PEI1* from S2 to S3 at 22°C in *toe1 toe2*. (E) Relative expression level of *TOE1* at 22°C and 16°C in Col-0 and *clf-28 swn-7*, as determined by qRT-PCR. Expression in Col-0 at 22°C was set as 1. Shown was mean ± SD of three biological replicates. Significance was calculated by Student’s *t*-test, ** *P* < 0.01; *** *P* < 0.001. (F) ChIP-qPCR assay showing enrichment of TOE1 at *LEC1*, *LEC2* and *PEI1* at S2 and S3 stages at 22°C and 16°C in Col-0. *AT4G05580* was a negative control. Data was mean ± SD of three replicates. Significance was calculated by Student’s *t*-test, * *P* < 0.05; ** *P* < 0.01; *** *P* < 0.001; ns, no significance. (G) A proposed working model for H2A.Z(ub) dynamics respond to temperature and H3K27me3 dynamics together determine cell fate during post-germination growth.

Given the reported association of TOE1 with PRC2 recruitment ^49^ and our confirmation of its interaction with CLF (Figures S7A and S7B), we also evaluated H3K27me3 level in *toe1 toe2* from S2 to S3 stages at 22°C. Unlike H2A.Z, in *toe1 toe2*, the increase of H3K27me3 level from S2 to S3 still occurred at the 341 genes (Figure S7C), including the cases such as *LEC1*, *LEC2* and *PEI1* (Figure S7D) This suggested that the deposition of H3K27me3 at embryonic genes from S2 to S3 stages may not be primarily mediated by TOE1. The dynamics of H2A.Z at the 341 genes in *toe1 toe2* at 22°C mimic those of Col-0 at 16°C (Figures S7E and S7F), implying a connection between TOE1 and the low-temperature response.

To investigate whether and how TOE1 responds to low temperature, we first measured the transcription level of *TOE1* at low temperature using RT-qPCR. The results revealed a decrease in *TOE1* expression at 16°C irrespective of the presence of PRC2 (Figure 7E). Next, we examined the binding of TOE1 to target genes at low temperature. Notably, there was reduced binding of TOE1 to chromatin regions of *LEC1*, *LEC2* and *PEI1* at 16°C compared to 22°C at both S2 and S3 stages (Figure 7F). These findings elucidate how the slow turnover of H2A.Z at embryonic related genes occurs at low ambient temperature.

These data suggest that TOE1 mediates INO80-C to evict H2A.Z at seed development and maturation-related genes, regulating the transition from embryonic to post-germination growth. Reduced temperatures reduced TOE1 expression and weaken the binding of TOE1 to targets, resulting in slower H2A.Z turnover. Such effect is independent of presence of PRC2-mediated H3K27me3. Consequently, the accumulation of H2A.Z(ub) at these genes rescues cell fate of *clf-28 swn-7* at low ambient temperature (Figure 7G).

## Discussion

Developmental plasticity enables plants to adapt to various environments. Ambient temperature, a common environmental cue, can induce changes in specific epigenetic modifications and developmental processes.^50–52^ In this study, using *prc2* mutant, we observed that low ambient temperature decelerates the turnover of H2A.Z at genes related to embryonic development during post-germination stages, regardless of PRC2 present. This accumulation of H2A.Z compensate for the absence of H3K27me3, thereby suppressing the ectopic expression of genes related to embryonic and seed development, thus promoting normal development. Furthermore, H2A.Z ubiquitination mediated by PRC1 and H3K27me3 mediated by PRC2 synergistically silence the seed development and maturation program, ensuring proper seed germination in Col-0.

### PRC1 catalyzes H2A.Zub to inhibit seed maturation program during germination

Previous research has established critical role of VAL-mediated PRC1 recruitment in suppressing seed maturation genes (*AFLs*) during germination, with PRC2 maintaining this repression.^14^ This conclusion is supported by genetic phenotypic evidence and the steady-state H2Aub and H3K27me3 profiles in various *prc1* and *prc2* mutants observed at *AFLs* genes in young seedling (10 DAS from the Yang et al., 2013) Here, we conducted a comprehensive analysis of H2Aub, H2A.Z, and H3K27me3 profiles globally, as well as H2A.Zub specifically on *AFLs* genes, at various stages of post-germinative growth: germination (S1), cotyledons fully opened (S2), and true leaves emerging (S3). This investigation aimed to elucidate the intricate interplay among different histone modifications in orchestrating the developmental program during post-germination phases.

Genome-wide, there were no significant changes in various histone modifications from S1 to S3. However, at embryonic and seed maturation genes, H3K27me3 level continuously increased, aligning with its function in maintaining repression post-germination. In contrast, H2Aub levels on the 341 genes remained unchanged across the three stages, contradicting with the previous finding that PRC1 initiate the inhibition of seed maturation program via H2Aub.^14^ Instead, H2A.Z level showed dynamic changes on these genes, increasing from S1 to S2, then declining at S3. Furthermore, a lack of H2A.Z led to higher expression of seed maturation related genes at S2 stage. In *clf-28 swn-7*, H2A.Z but not H2A, function in repression of seed maturation genes, restoring polycomb callus at 16°C. These results indicate H2A.Z play roles during seed germination, consistent with the previous study showing delayed seed germination in *hta9hta11* and *pie1-5* mutants, which impair H2A.Z deposition.^25^ Additionally, the morphology of *hta9-1 clf-28 swn-7* resembles *atbmi1a/b/c* mutants before cotyledons fully opened. H2A.Z could also be monoubiquitinated by PRC1 to repress gene expression.^9^ ChIP-re-ChIP-qPCR assays confirmed a similar pattern of H2A.Zub as H2A.Z on *AFLs* genes from S1 to S3 stages. Thus, H2A.Zub, catalyzed by PRC1, plays a role in initiating the repression of seed maturation programs during germination. Understanding how PRC1 selectively catalyzes H2Aub or H2A.Zub on different targets during specific development programs or in response to environmental cues remains an intriguing area for further research.

### Low temperature selectively slows down H2A.Z turn-over via TOE1

Previous studies have shown that H2A.Z responds to temperature fluctuations: it restricts chromatin accessibility and suppresses transcriptome activity under low temperatures,^26^ while being rapidly removed from temperature-responsive genes at higher temperatures.^33^ Here, we focus on the prolonged exposure to low temperature, revealing dynamic changes in H2A.Z levels. Under 22°C, H2A.Z levels declined specifically at 341 genes related to embryonic and seed maturation from stage S2 to S3. Conversely, at 16°C, H2A.Z remained elevated at these genes across the same stages, likely due to a slowdown in turnover rates. The regulation of H2A.Z turnover involves INO80-C, which is targeted by specific transcription factors.^30,53,54^ Our findings indicate that TOE1, enriched at the binding sites of these 341 genes, binds to key targets such as *LEC1* and *LEC2*, as confirmed by ChIP-qPCR, and physically interacts with the INO80-C components EED and ARP5. Loss of TOE1 leads to high accumulation of H2A.Z at 22°C from stage S2 to S3. Moreover, transcription of TOE1 is reduced at 16°C, further contributing to decreased TOE1 binding at target genes. Collectively, these factors contribute to the selectively slowed turnover of H2A.Z at embryonic and seed maturation genes under 16°C. Notably, low temperature dramatically reduce PRC2 binding to targets, resulting in a slight reduction in H3K27me3 levels, yet gene expression remains unchanged likely due to compensatory prolonged abundance of H2A.Z. Additionally, the selective accumulation of H2A.Z at H3K27me3 targets induced by low temperature occurs in both *prc2* mutants and Col-0, suggesting a conserved mechanism independent of PRC2. Further investigation is required to fully comprehend how ambient temperature influences the interplay between transcription factors and chromatin regulators.

### Low temperature modulates chromatin modification interplay to change cell fate

Our study unveils that H2A.Z compensates for the absence of H3K27me3 at cell fate-related genes under low temperature in *prc2* mutants, highlighting the synergistic role of epigenetics and environmental conditions in determining cell fate. It has been reported that H2A.Z and H3K27me3 are highly correlated at poised enhancers^24^ and 90% of H3K27me3 is co-located with H2A.Z in *Arabidopsis*.^55^ Therefore, the compensation effect of H2A.Z for loss of H3K27me3 is logical. Additionally in mouse embryonic stem cells, H2A.Z facilitates PRC2 activity for H3K27me3 by promoting chromatin compaction.^56^ Thus, it’s possible that H2A.Z facilitates the accumulation of H3K27me3 after germination in *Arabidopsis*.

The temperature-triggered alteration of cell fate occurs at early post-germination stages, emphasizing the importance of the developmental window. Slower growth due to low temperature may extend the developmental window on determination of cell fate. This ambient temperature-induced interplay of H2A.Z and H3K27me3 mechanism may be borrowed by other eukaryotes. In humans, H3K27me3 dysregulation has been found in various cancers. For instance, H3K27me3 loss is an independent risk factor for meningioma recurrence.^57^ Understanding the mechanisms of low temperature-induced epigenetic compensation could inform cancer treatments. By targeting transcription factors that play a recruiting role, it may be possible to induce the accumulation of H2A.Z or other inhibitory modifications to compensate for the loss of H3K27me3, potentially inhibiting cancer cell proliferation.

In summary, we have uncovered that low temperature influences cell developmental fate by modulating the dynamics of H2A.Z and H3K27me3. In Col-0, H2A.Zub and H3K27me3 cooperatively initiate and maintain repression of seed maturation program, ensuring proper seed germination and seedling development. In *prc2* mutants, the absence of H3K27me3 fails to sustain the repression of embryonic and seed maturation genes, leading to callus formation instead of true leaves. However, under low temperature, the slow turnover rate of H2A.Z, likely due to impaired TOE1’s recruitment of INO80-C, allows H2A.Z to persist on embryonic-related genes, compensating for the lack of H3K27me3 repression after cotyledons fully open. Consequently, the likelihood of ‘plant’ architecture increases under low temperature (Figure 7G).

### Limitations of the study

First, although our research has showed that H2A.Zub rather than H2Aub mediated by PRC1 initiates inhibition of seed maturation program during post-germination development stage, the mechanism by which PRC1 selectively catalyzes H2Aub or H2A.Zub on different targets during specific development programs or in response to environmental cues remains unknown. Second, we confirmed the interaction between TOE1 with both INO80-C and PRC2. However, whether TOE1 recruits these complexes and regulate H2A.Z and H3K27me3 dynamics simultaneously during specific biological processes remains an intriguing area for further research. Third, the response of TOE1 to low temperature deserves further study.

## STAR*Methods

### Key Resources Table

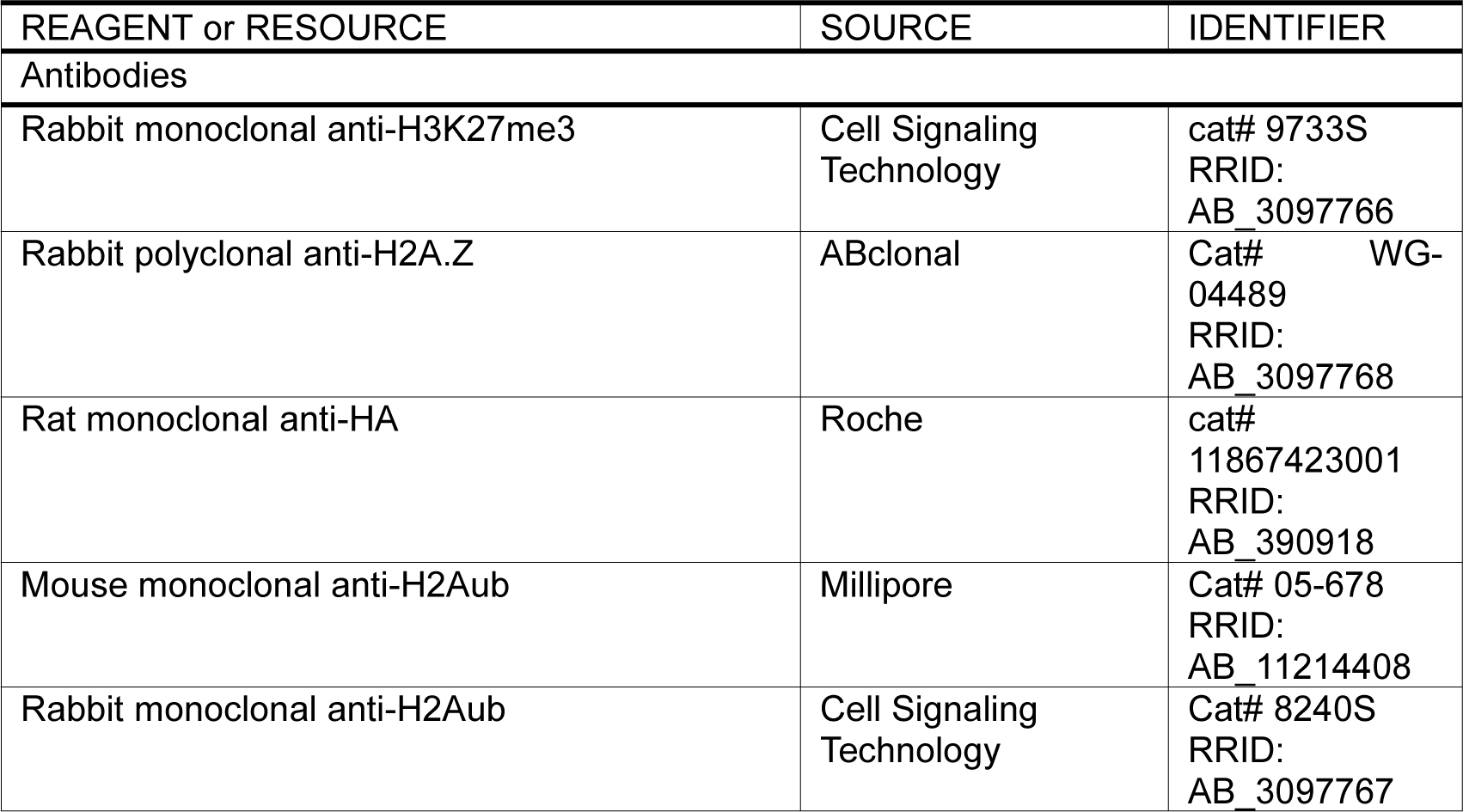
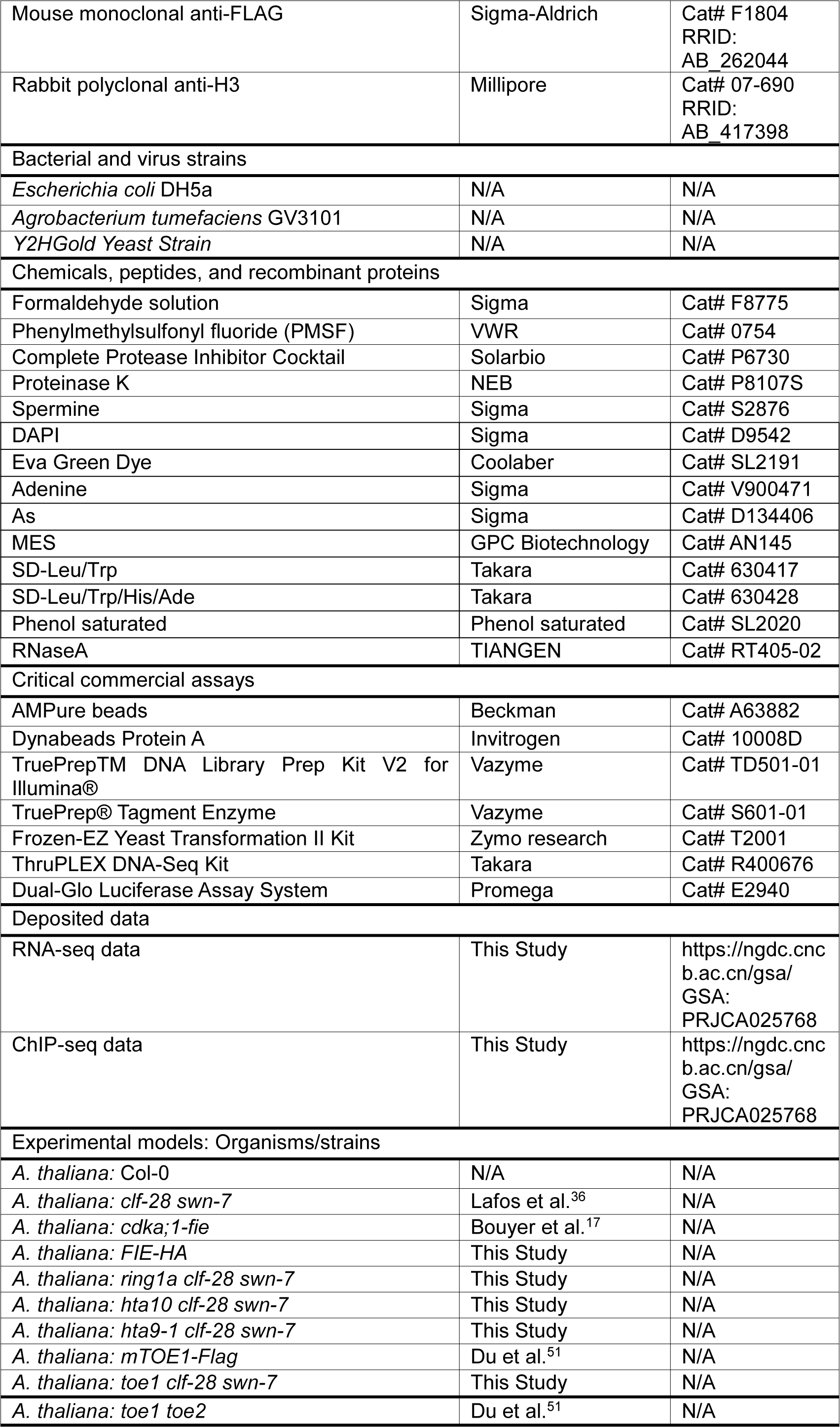
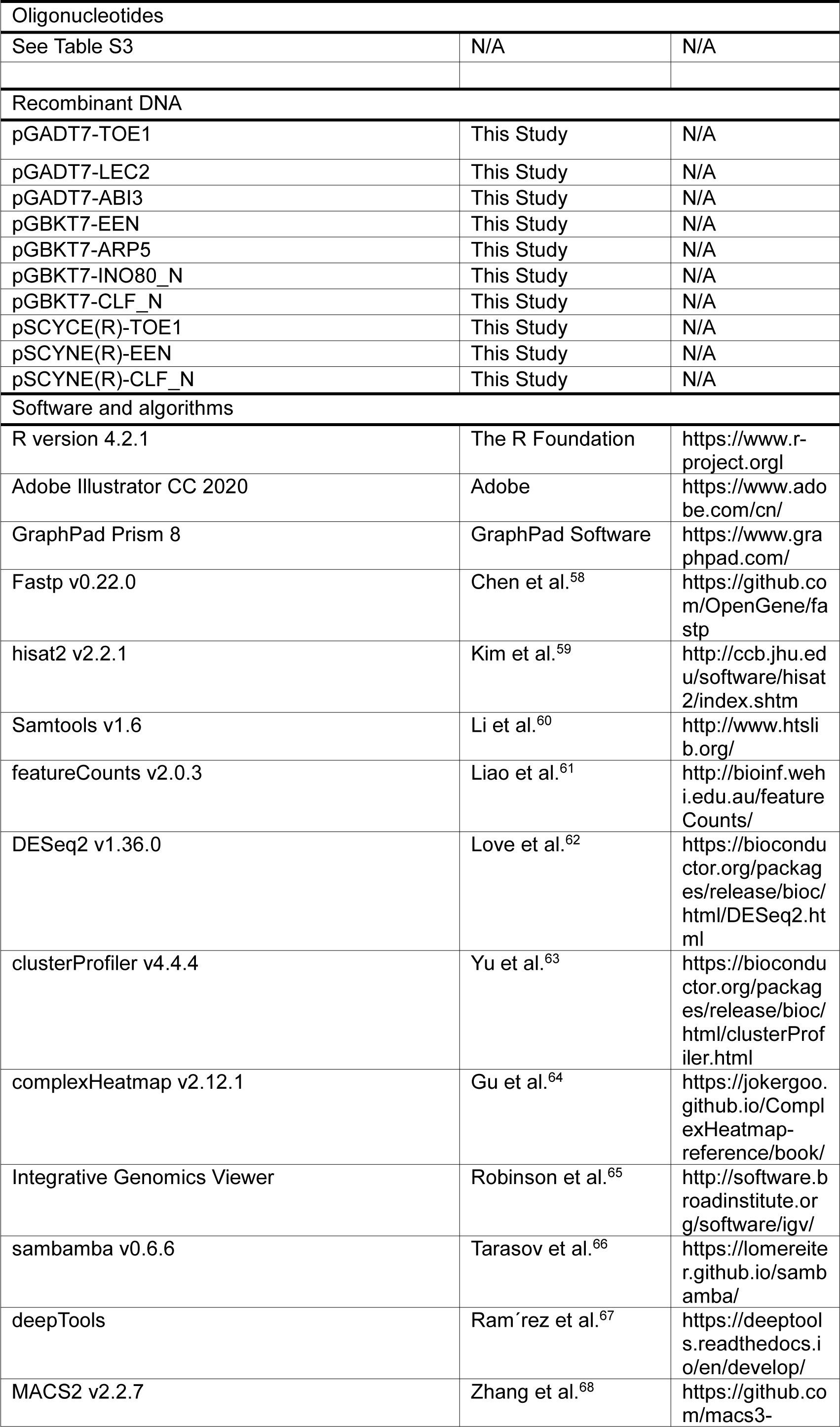
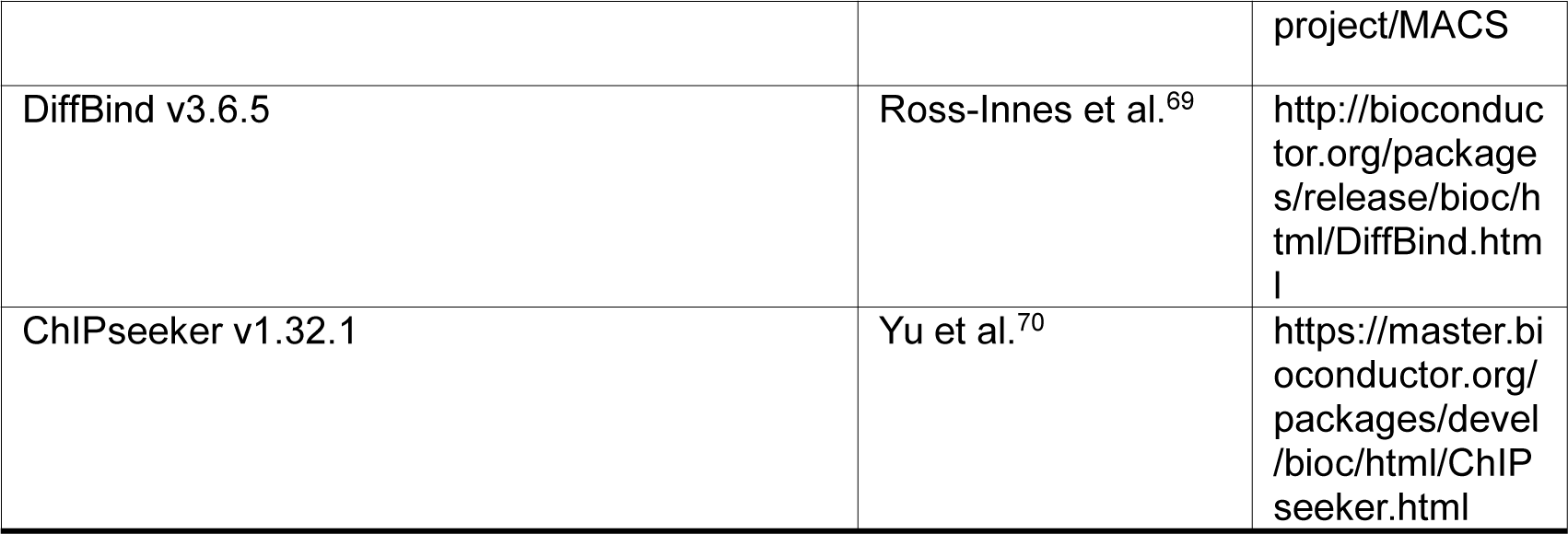

### Resource Availability

#### Lead Contact

Further information and requests for resources should be directed to and will be fulfilled by the Lead Contact, Jun Xiao.

### Materials Availability

Plasmids and genetic materials generated in this study will be made available on request to the lead contact. This study did not generate new unique reagents.

### Data and Code Availability

The ChIP-seq and RNA-seq data (BioProject PRJCA025768) were deposited in Beijing Institute of Genomics Data Center (http://bigd.big.ac.cn).

Code used for all processing and analysis is available at GitHub (https://github.com/kh-zhu/prc2-h2az-temp).

## EXPERIMENTAL MODEL AND SUBJECT DETAILS

### Plant materials and growth conditions

The *Arabidopsis thaliana* plants were used as the experimental model in the study. *Arabidopsis thaliana* accession Col-0 was used as the background in all experiments. The double mutant *clf-28 swn-7* and *cdka;1-fie* have been described previously.^17,36^ The triple mutant *ring1a clf-28 swn-7, hta9-1 clf-28 swn-7* and *hta10 clf-28 swn-7* were generated by crossing *clf-28 swn-7* with *ring1a, hta9-1* and *hta10* respectively. The *mTOE1-Flag* has been described previously.^48^ Besides, *toe1 clf-28 swn-7* triple mutant were generated by crossing *toe1 toe2* and *clf-28 swn-7*.^48^ The seeds were sown on 1/2 Murashige and Skoog medium with 1% sucrose and 0.8% agar and stratified at 4°C for 3 days. Then, the seeds were transferred into 22°C and 16°C long day (16 hours light and 8 hours dark) conditions respectively.

### Method Details

#### Imaging of microscopy observation

The phenotype of mutants *clf-28 swn-7*, *ring1a clf-28 swn-7*, *hta9-1 clf-28 swn-7*, *hta10 clf-28 swn-7* and *toe1 clf-28 swn-7* at 22°C and 16°C were imaged using a Zeiss microscope (SteREO Discovery.V12).

#### RNA extraction, qRT-PCR analysis, and RNA-seq

For Col-0, seedling grown under long-day conditions for 4 days and 6 days at 22°C, 6 days and 8 days at 16°C, which correspond to S3 and S4 stages respectively. For *clf-28 swn-7*, seedlings were selected from S2 to S5, the time were displayed at Figure 2A. For *hta9-1 clf-28 swn-7*, 14 days/14 days at 22°C/16°C. For *toe1 clf-28 swn-7*, 35 days/35 days at 22°C/16°C. Total RNA was extracted using Quick RNA Isolation Kit with on-column DNaseI digestion (Huayueyang, Cat# 0416-50), according to the manufacturer’s protocol. Each line and condition set three independent biological replicates. For qRT-PCR, first-strand cDNA was reverse transcribed using FastKing RT kit (TIANGEN, Cat# KR116). Subsequent qRT-PCR assays were performed using the SYBR qPCR Master Mix (Vazyme Biotech, Q711-03), relevant primer sequences are given in Table S3. *Actin 2* (*AT3G18780*) was used to normalize expression levels. For RNA-seq, library construction and sequencing were performed by the Illumina NovaSeq platform (Annoroad, Beijing, China).

#### Western blotting

Protein samples were extracted from seedlings growing for 7 days and separated on SDS-PAGE gels, followed by transferred to PVDF membranes. After blocking with 5% milk in PBST (0.1% Tween-20) for 1 h, the membrane was incubated with anti-H2Aub (Millipore, Cat# 05-678) and anti-H3 (Millipore, Cat# 07-690) for 2 h at room temperature. Then, membranes were washed using PBST and incubated with secondary antibodies for 1 h at room temperature. Finally, membranes were washed using PBST and the signals were examined by a chemiluminescent HRP substrate (EASYBIO, Cat# BE6706).

#### ChIP-qPCR and ChIP-seq Assays

For ChIP experiment, *clf-28 swn-7*, Col-0, mTOE1-Flag and *toe1 toe2* for different stages and different temperatures were harvested and fixed according to a published protocol.^71^ The chromatin extract was immunoprecipitated with specific histone marker antibody. The antibodies used for immunoprecipitate are anti-H3K27me3 (Cell Signaling Technology, Cat# 9733S), anti-H2A.Z (ABclonal, Cat# WG-04489), anti-H2Aub (Cell Signaling Technology, Cat #8240S), and anti-Flag (Sigma-Aldrich, Cat# F1804). The immunoprecipitated DNAs were incubate with mixed protein A Dynabeads (Invitrogen, Cat# 10008D). ChIP DNAs were reverse crosslinked and was extracted with phenol to chloroform to isoamyl alcohol (25:24:1), precipitated with ethanol and resuspended in ddH2O. For ChIP-qPCR, the amounts of immunoprecipitated DNA of H3K27me3 and H2A.Z were normalized to *SEP3* and *TUB2* respectively, the enrichment of TOE1 was normalized to background signal (Fold enrichment). All primer information was list in Table S3. For ChIP-seq, libraries were amplified using ThruPLEX DNA-Seq Kit (Takara, Cat# R400676) according to the manual and purified by AMPure XP beads (Beckman, Cat# A63882). The libraries were sequenced using an Illumina Novaseq platform by Annoroad Gene Technology. Two biological replicates were performed.

#### CUT&Tag-qPCR experiments

The CUT&Tag experiment was performed using *clf-28 swn-7* at S4 stage (13 DAS at 22°C and 15 DAS at 16°C). The CUT&Tag experiment was performed according to previously described protocols,^72^ with a minor modification: each sample contained apporximately 1 million nuclei. For qPCR ^47^, half of the DNA products of CUT&Tag was used as “Input”, and the other half of DNA undergoing library PCR amplification and purification was used as “IP products”.^47^ The “Input” and “IP products” were dilute 30 times for qPCR assay (Equivalent to 100% Input). The qPCR reaction system and procedure are the same as ChIP-qPCR. The fold enrichment of H2A.Z was normalized to the amount of input chromatin, which was calculated in the same way as ChIP-qPCR. The fold enrichment value >1 indicates enrichment of binding sites.

The primers for specific regions are provided in in Table S3.

#### ChIP-Re-ChIP-qPCR

The ChIP-re-ChIP experiment was performed using Col-0 at S1-S3 stage according to a published protocol.^73^ The procedure were same as ChIP before elution the ChIP DNA from protein A beads (Figure S5B). The anti-H2A.Z (ABclonal, Cat# WG-04489) was used for first immunoprecipitation. After first elution, anti-H2A.Z, anti-H2Aub (Millipore, Cat# 05-678) and no antibody were used for second immunoprecipitation. The following incubation, reverse crosslink and re-ChIP DNA extraction were same as ChIP. For re-ChIP-qPCR, the fold enrichment of H2A.Z and H2A.Zub was normalized to background signal (H2A.Z-no Antibody). All primer information was list in Table S3.

#### Yeast two-hybrid assay

Yeast two-hybrid assays were performed according to the manual of Frozen-EZ Yeast Transformation II™ kit (Zymo Research, Cat# T2001). For the interaction, the CDS fragments of *TOE1*, *LEC2*, *ABI3* were cloned into the prey vector (pGADT7), and the fragments of *EEN*, *APR5*, *INO80_N*, *CLF_N* into the bait vector (pGBKT7) vector. The cassettes plasmids were transformed into yeast strain Y2HGold. Transformed yeast cells were grown on either SD-Trp/Leu and SD-Trp/Leu/His/Ade plates at 30°C for 3-4 days. The primers are listed in Table S3.

#### Bimolecular fluorescence complementation (BiFC) assay

The vectors used to make constructs for BiFC vectors were pSCYNE(R) (nCFP) and pSCYCE(R) (cCFP), which carry fragments encoding the N- and C-terminal of CFP, respectively. For the interaction, the CDS fragments of *TOE1* was cloned into pSCYCE(R), and the fragments of *EEN* and *CLF_N* were cloned into pSCYNE(R). Young leaves of 4-week-old *N. benthamiana* plants were co-infiltrated with *A. tumefaciens* strain GV3101 harbouring different combinations of these plasmids, and the GV3101 cultures harboring constructs expressing cCFP fusion proteins and nCFP fusion proteins were mixed at a ratio of 1:1. *N. benthamiana* plants grown in long-day conditions for 48 h-72 h after infiltration, the CFP signals were detected by a confocal microscope (LSM980; Carl Zeiss, Germany). H2B-mCherry was used as a cell nucleus marker. The primers are listed in Table S3.

### Quantification and Statistical Analysis

#### RNA-seq data analyses

For each library, raw.fastq was performed quality control and filtering using fastp v0.22.0 with default parameters.^58^ The trimmed reads were aligned to the Arabidopsis genome (TAIR10) using hisat2 v2.2.1.^59^ The resulting SAM file was converted into BAM format, sorted and indexed with samtools v1.6.^60^ Paired mapping reads overlapping each annotated gene (Araport11) were counted using featureCounts v2.0.3.^61^ The counts files were normalized by Transcripts Per Kilobase of exon model per Million mapped reads (TPM). Differential gene expression analysis was performed using DESeq2 v1.36.0,^62^ with the threshold for differential gene expression set at “*p.adjust* < 0.05 and abs(log2FoldChange) >1”. The PCA plot was generated by FactoMineR v2.10 package of R.^74^ Heatmap were generated by “k-means” function in R and ComplexHeatmap v2.12.1 package.^64^ The GO enrichment analysis of differentially expressed genes were conducted using clusterProfiler v4.4.4.^63^ The motif scanning was performed by fimo v5.4.1.^75^

#### ChIP-seq data analyses

For each library, raw.fastq was also performed quality control and filtering using fastp v0.22.0 with default parameters. Reads were aligned to the Arabidopsis genome (TAIR10) using bwa v0.7.17.^76^ The resulting SAM file was converted into BAM format and sorted using samtools v1.6.^60^ Duplicated reads were removed using sambamba v0.6.6 and index files were created by samtools v1.6.^66^ Two biological replicates were merged with samtools v1.6. To normalize and visualize the individual and merged replicate datasets, the BAM files were converted to bigwig using bamCoverage v3.5.1 provided by deepTools with a bin size of 10 bp and normalized by Counts Per Million (CPM).^67^ The “plotProfile” and “PlotHeatmap” functions in deepTools were used to check the quality of samples, and scored associated genomic regions. All the signal plots were visualized by Integrative Genomics Viewer.^65^ Macs2 v2.2.7.1 was used to call peaks,^68^ with the “broad” parameter for the H3K27me3 ChIP-seq datasets analysis. DiffBind v3.6.5 was used to quantity peaks value with following parameters “ score = DBA_SCORE_TMM_READS_EFFECTIVE_CPM, bUseSummarizeOverlaps = TRUE, bParallel = TRUE, summits = T “,^69^ and DESeq2 was used to identify differential binding peaks. The peaks called by macs2 and differential peaks identified by DiffBind v3.6.5 were annotated using TxDb.Athaliana.BioMart.plantsmart28 and ChIPseeker v1.32.1 package with “annotatePeak” function with the parameter “tssRegion = c(−1500, 500)”.^70^ The GO analysis for ChIP-seq datasets was performed following the methods used for RNA-seq analysis.

## Acknowledgments

We thank Arabidopsis Biological Resource Center for T-DNA insertion lines; Dr. Danhua Jiang (Institute of Genetics and Developmental Biology, Chinese Academy of Sciences) for the h2a.z seeds; Dr. Hongquan Yang (Shanghai Key Laboratory of Plant Molecular Sciences, College of Life Sciences, Shanghai Normal University) for the mTOE1-Flag and toe1toe2 seeds; members in J.X. lab for discussion and comments on the manuscript. This work was supported by the National Key Research and Development Program of China (2021YFD1201500), Beijing Natural Science Foundation Outstanding Youth Project (JQ23026), the CAS Project for Young Scientists in Basic Research (YSBR-093) to J.X., and National Science Foundation grant MCB 2224729 to D.W.

## Author Contributions

J.X. designed the research; D.W. provided some input into research design and manuscript drafting; K.-H.Z. performed most of the experiments with help from F.-F.L.; L.Z. and K.-H.Z. analyzed ChIP-seq and RNA-seq data; X.-L. L, C.-S.H. and D.W. commented and polished the manuscript; K.-H.Z. and J.X. wrote the manuscript.

## Supplemental information

Document S1. Figures S1–S7

Table S1. RNA-seq data used in this study.

Table S2. ChIP-seq data used in this study.

Table S3. List of primers used in this study, related to STAR Methods

---

**Figure S1.**
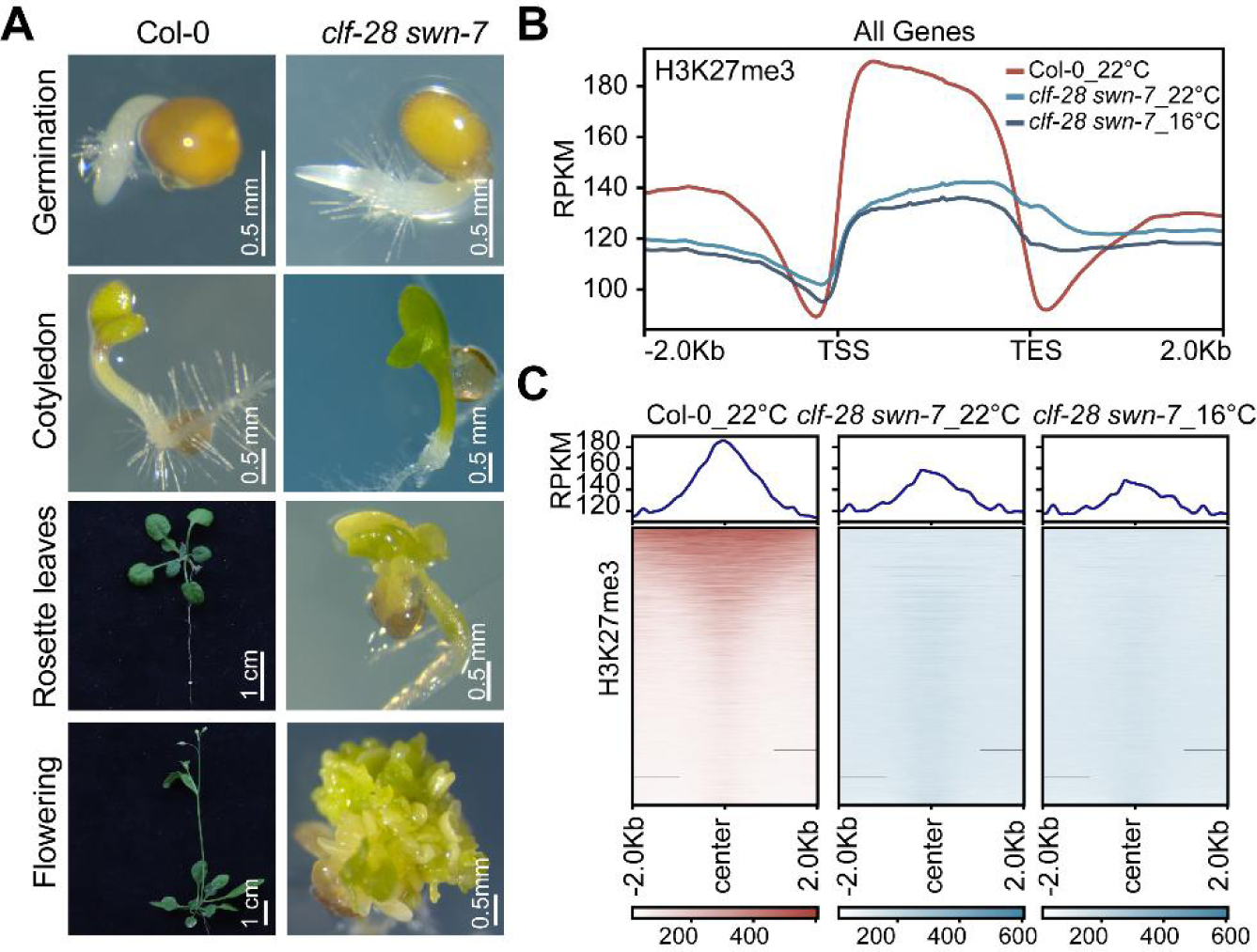
Developmental defects and H3K27me3 profile of *clf-28 swn-7*. (A) The developmental defects of *clf-28 swn-7* compared to Col-0 at 22°C at different stages. Scale bar is as indicated. (B and C) ChIP-seq profile of H3K27me3 along genes (B) or peak summit (C) showing the dramatic reduction of H3K27me3 in *clf-28 swn-7* compared to Col-0 at 22°C, not restored at 16°C.

**Figure S2.**
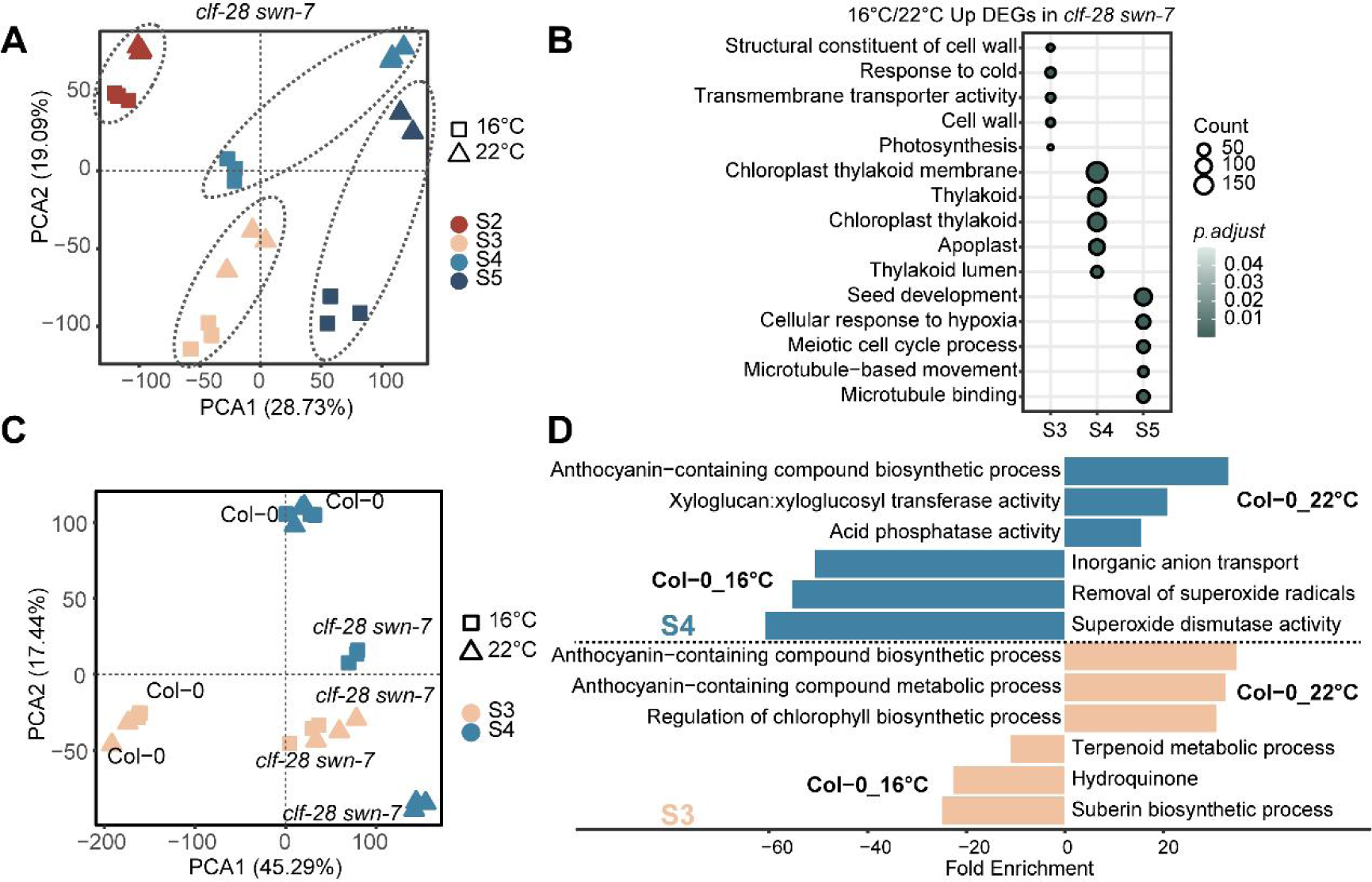
PCA analysis and GO enrichment of DEGs for Col-0 and *clf-28 swn-7* at different developmental stages. (A) PCA analysis of transcriptomes in *clf-28 swn-7* at S2 to S5 stages under two ambient temperatures. (B) GO enrichment analysis of up-regulated genes in *clf-28 swn-7* at 16°C compared to 22°C from S3 to S5 stages. (C) PCA analysis of transcriptomes in Col-0 and *clf-28 swn-7* at S3 and S4 stages under two ambient temperatures. (D) GO enrichment analysis of genes with elevated expression under either 16°C or 22°C at S3 and S4 stages in Col-0.

**Figure S3.**
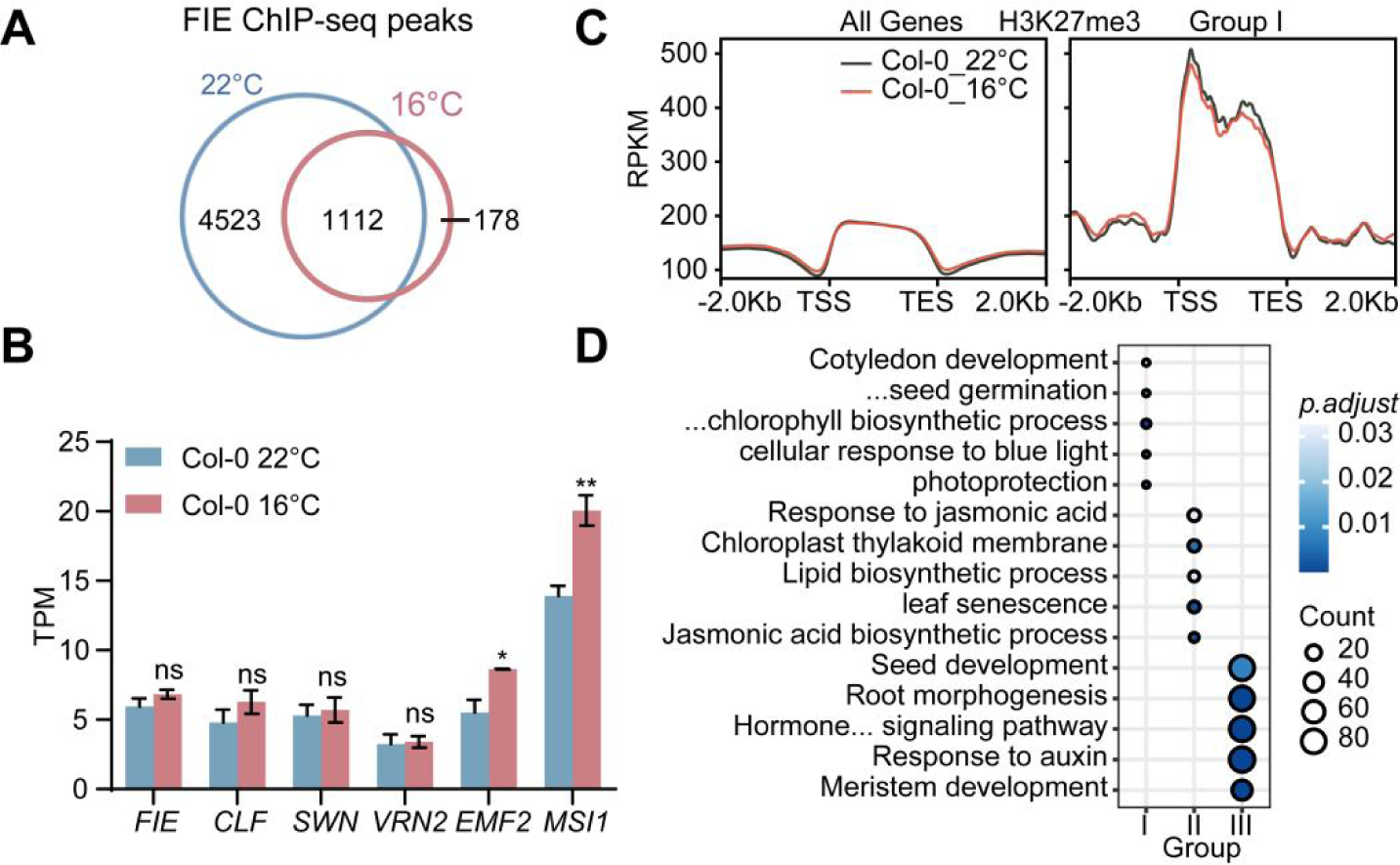
Overview of FIE direct targets and H3K27me3 profile in Col-0 at different temperatures. (A) Venn diagram showing number of FIE binding peaks at 22°C and 16°C. (B) Expression changes of PRC2 components coding genes between 22°C and 16°C in Col-0 at S3. Data was mean ± SEM of three biological replicates from RNA-seq. Student’s *t*-test, * *P* < 0.05; ** *P* < 0.01; ns, no significant changes. (C) ChIP-seq profile of H3K27me3 along genic region of all genes or abolished FIE-binding and up-regulated genes (Group I, Figure 3D) at 22°C and 16°C in Col-0. TSS, transcription start site; TES, transcription end site. (D) GO enrichment analysis of three group of genes from Figure 3D.

**Figure S4.**
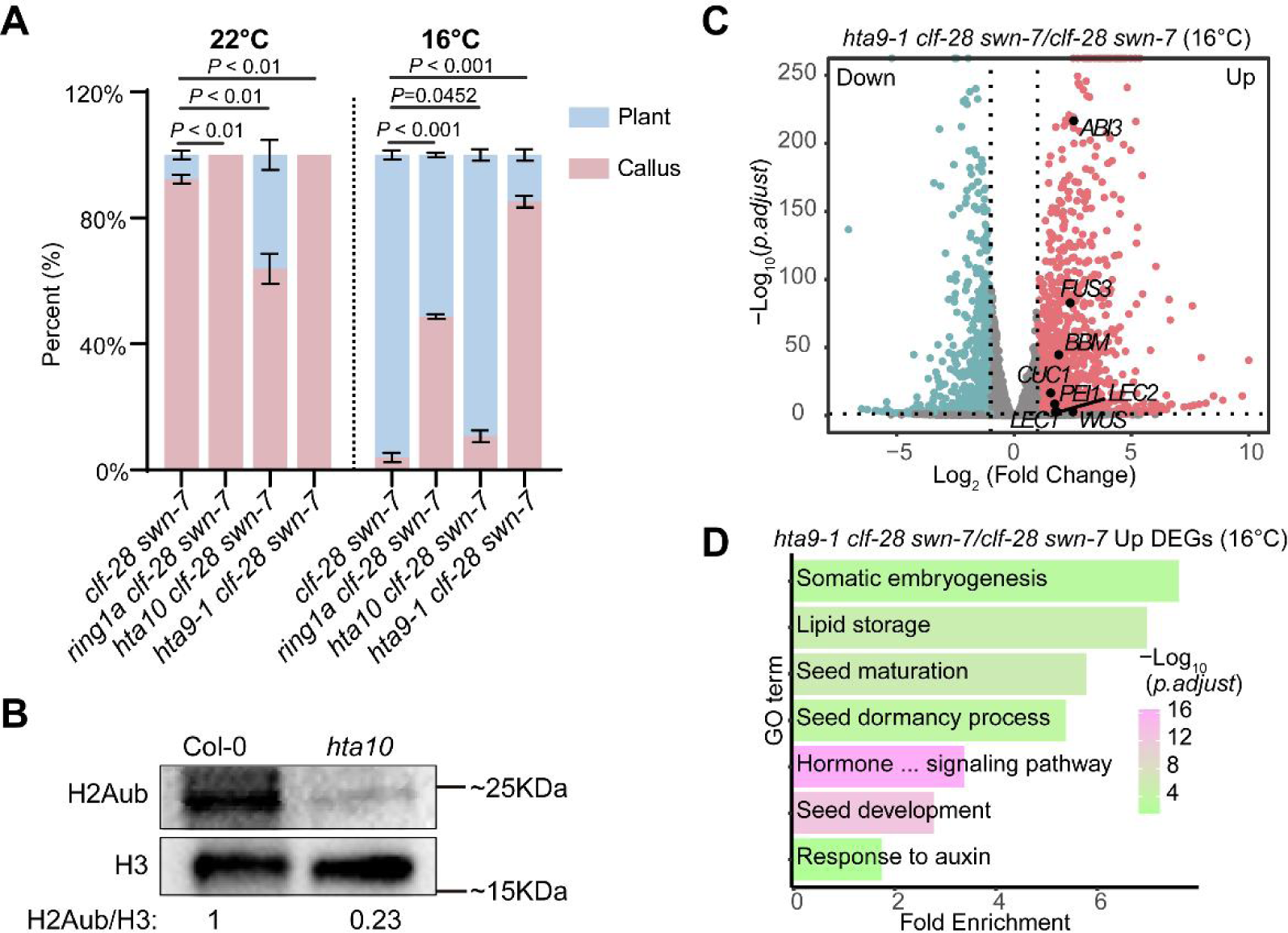
The various ‘callus’ and ‘plant’ proportions of different high order mutants. (A) The various proportion of ‘callus’ and ‘plant’ in *clf-28 swn-7*, *ring1a clf-28 swn-7*, *hta10 clf-28 swn-7* and *hta9-1 clf-28 swn-7* at 22°C and 16°C. Data was mean ± SEM from three biological replicates, n >= 30 for each replicate. Significance was calculated by Student’s *t*-test. (B) Western blot assay showing H2Aub levels in Col-0 and *hta10* mutant. (C) Volcano plot of genes with lower (blue) or higher (pink) expression in *hta9-1 clf-28 swn-7* compared to *clf-28 swn-7* at 16°C. Gray dots, no difference between two samples. The known genes were labeled and shown in black dots. The cutoff of significance: Log_2_ (Fold change) > 1 or < −1; *p.adjust* < 0.05. (D) GO enrichment analysis of the up-regulated genes in *hta9-1 clf-28 swn-7* compared to *clf-28 swn-7* at 16°C.

**Figure S5.**
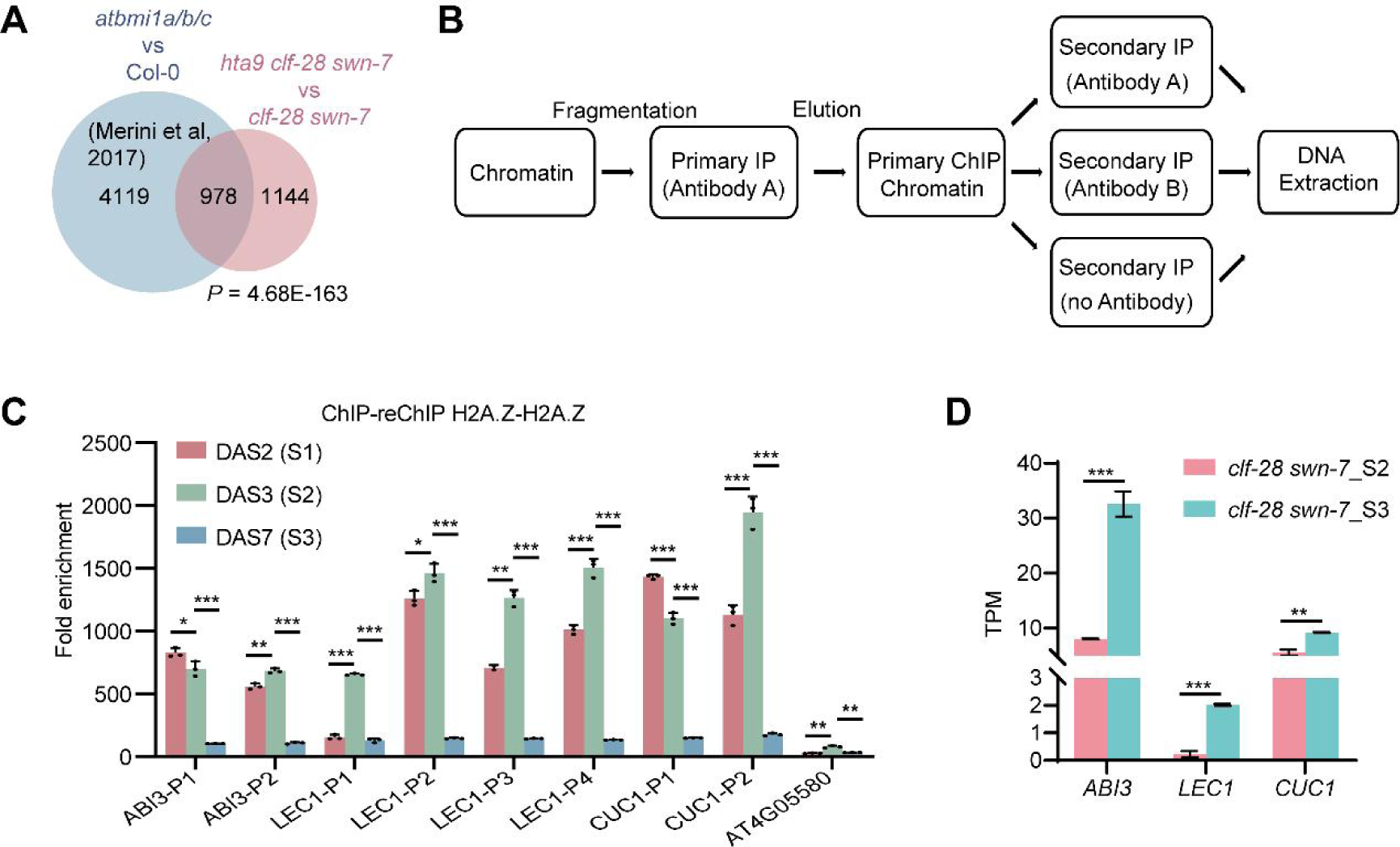
H2A.Zub and H3K27me3 coordinately participate in silencing seed development genes during post-germination. (A) Venn diagram showing significant overlap of DEGs between *atbmia/b/c* and Col-0 with DEGs between *hta9-1 clf-28 swn-7* and *clf-28 swn-7*. Fisher exact-test, *P* = 4.68E-163. (B) Diagram of re-ChIP assay. (C) ChIP re-ChIP qPCR showing H2A.Z pattern at *ABI3*, *LEC1* and *CUC1* loci from 2 DAS (S1) to 7 DAS (S3) in Col-0 at 22°C. *AT4G05580* as a negative control. Data was mean ± SD of three repeats. (D) Expression level of A*BI3*, *LEC1* and *CUC1* from S2 to S3 in *clf-28 swn-7* mutant at 22°C. Data was mean ± SD of three biological replicates from RNA-seq. Student’s *t*-test, ** *P* < 0.01; *** *P* < 0.001.

**Figure S6.**
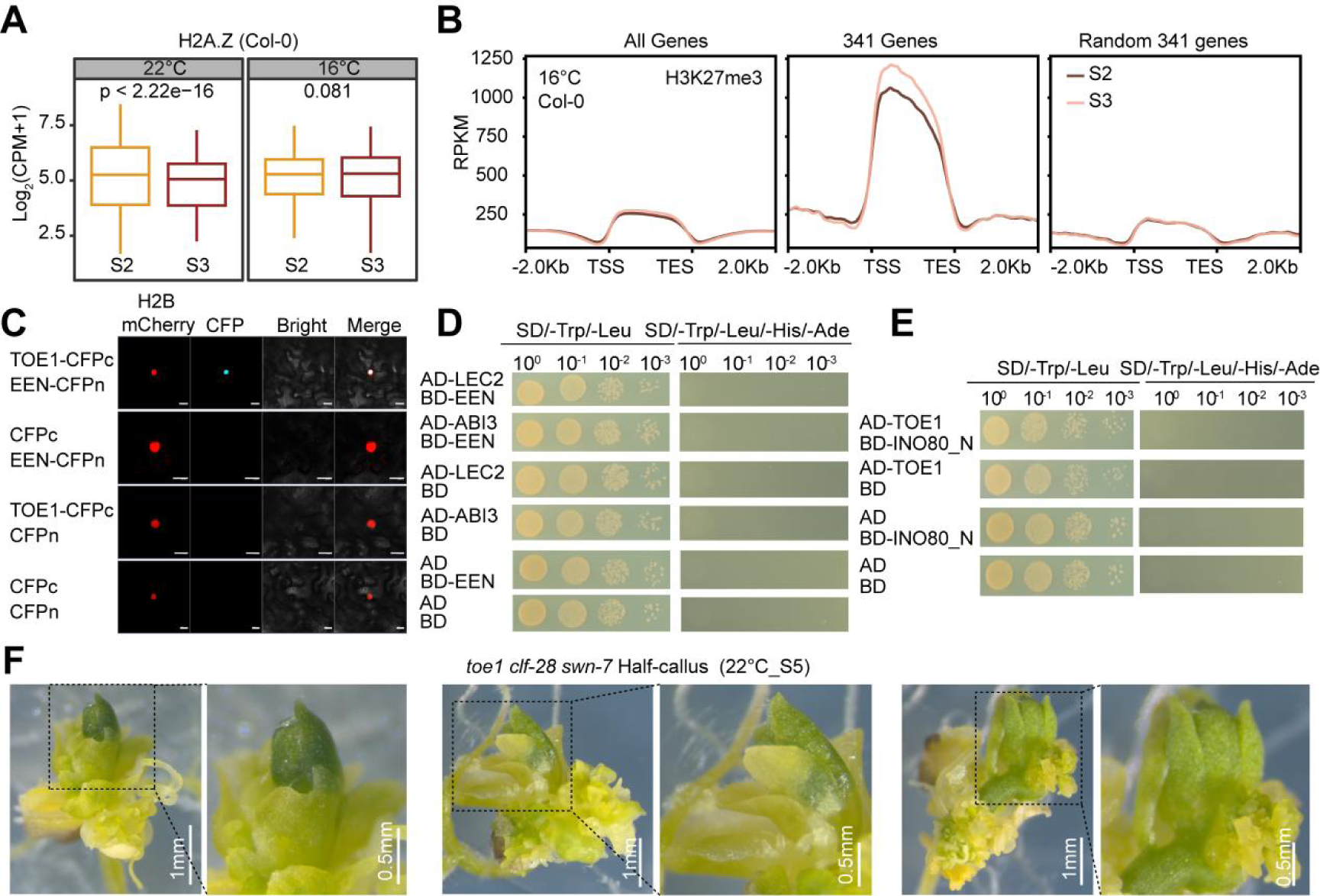
Interactions of TOE1 with other factors. (A) Boxplot showing level of H2A.Z peaks along the 341 genes (Figure 3F) between S2 and S3 at 16°C and 22°C in Col-0, respectively. Student’s *t*-test are performed. (B) ChIP-seq profile of H2A.Z and H3K27me3 along genic region of all genes, the 341 genes (Figure 3F) and the other random 341 genes from S2 to S3 at 16°C in Col-0. TSS, transcription start site; TES, transcription end site. (C) BiFC assay showing the interaction between TOE1 and INO80-C (EEN). H2B-mCherry is a nuclear maker. Scale bar, 20 µm (D) Y2H assay showing interaction results of TOE1 with other transcription factors (Figure 6E). (E) Y2H assay showing interaction results of TOE1 with other component (INO80) of INO80-C. (F) ‘Half-callus’ morphology of *toe1 clf-28 swn-7*, pictures were taken for 35 days at 22°C. Scale bar is as indicated.

**Figure S7.**
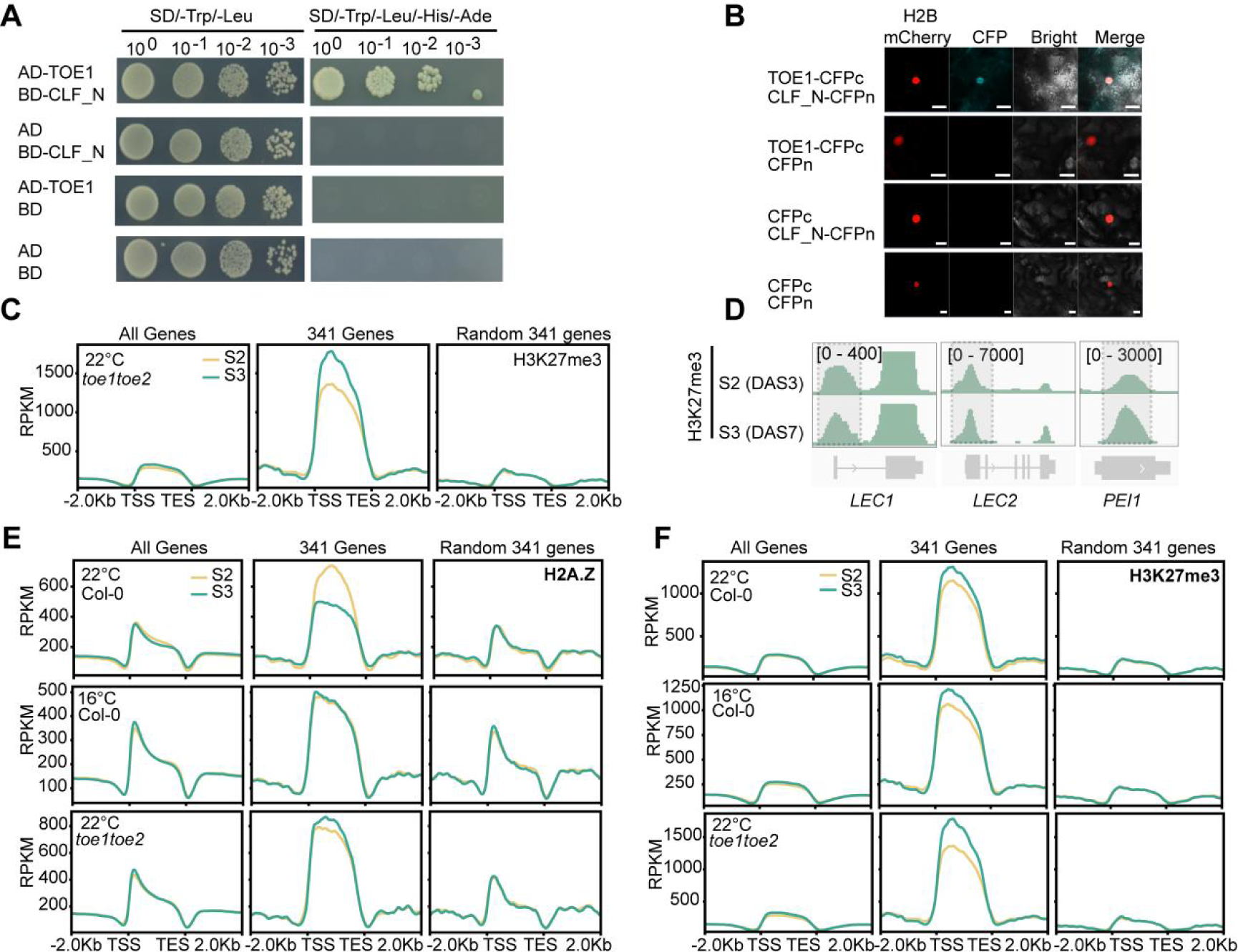
H3K27me3 dynamics in *toe1 toe2* at 22°C. (A) Y2H assay showing the interaction between TOE1 and CLF. (B) BiFC assay showing the interaction between TOE1 and CLF. H2B-mCherry is a nuclear maker. Scale bar, 20 µm. (C) ChIP-seq profile of H3K27me3 along genic region of all genes, the 341 genes (Figure 3F) and the other random 341 genes from S2 to S3 at 22°C in *toe1 toe2*. TSS, transcription start site; TES, transcription end site. (D) IGV browser view of H3K27me3 signal at *LEC1*, *LEC2* and *PEI1* from S2 to S3 at 22°C in Col-0 and *toe1 toe2*. (E and F) ChIP-seq profile of H2A.Z (E) and H3K27me3 (F) along genic region of all genes, the 341 genes (Figure 3F) and the other random 341 genes from S2 to S3 at 22°C in Col-0, 16°C in Col-0, 22°C in *toe1 toe2*. TSS, transcription start site; TES, transcription end site.

